# Chromatin accessibility dynamics of *Chlamydia*-infected epithelial cells

**DOI:** 10.1101/681999

**Authors:** Regan J. Hayward, James W. Marsh, Michael S. Humphrys, Wilhelmina M. Huston, Garry S.A. Myers

**Affiliations:** The ithree institute, University of Technology Sydney, Australia; Max Planck Institute for Developmental Biology, Tuebingen, Germany; Institute for Genome Sciences, University of Maryland School of Medicine, Baltimore, MD United States; School of Life Sciences, Faculty of Science, University of Technology Sydney, Australia

**Keywords:** Chlamydial infection, *Chlamydia trachomatis*, chromatin accessibility, Faire-Seq, bacterial infection

## Abstract

*Chlamydia* are Gram-negative, obligate intracellular bacterial pathogens responsible for a broad spectrum of human and animal diseases. In humans, *Chlamydia trachomatis* is the most prevalent bacterial sexually transmitted infection worldwide and is the causative agent of trachoma (infectious blindness) in disadvantaged populations. Over the course of its developmental cycle, *Chlamydia* extensively remodels its intracellular niche and parasitises the host cell for nutrients, with substantial resulting changes to the host cell transcriptome and proteome. However, little information is available on the impact of chlamydial infection on the host cell epigenome and global gene regulation. Regions of open eukaryotic chromatin correspond to nucleosome-depleted regions, which in turn are associated with regulatory functions and transcription factor binding. We applied Formaldehyde-Assisted Isolation of Regulatory Elements enrichment followed by sequencing (FAIRE-Seq) to generate temporal chromatin maps of *C. trachomatis*-infected human epithelial cells *in vitro* over the chlamydial developmental cycle. We detected both conserved and distinct temporal changes to genome-wide chromatin accessibility associated with *C. trachomatis* infection. The observed differentially accessible chromatin regions, including several Clusters of Open Regulatory Elements (COREs) and temporally-enriched sets of transcription factors, may help shape the host cell response to infection. These regions and motifs were linked to genomic features and genes associated with immune responses, re-direction of host cell nutrients, intracellular signaling, cell-cell adhesion, extracellular matrix, metabolism and apoptosis. This work provides another perspective to the complex response to chlamydial infection, and will inform further studies of transcriptional regulation and the epigenome in *Chlamydia*-infected human cells and tissues

## Introduction

Members of the genus *Chlamydia* are Gram-negative, obligate intracellular bacterial pathogens responsible for a broad spectrum of human and animal diseases [1]. In humans, *Chlamydia trachomatis* is the most prevalent bacterial sexually transmitted infection (STI) [2], causing substantial reproductive tract disease globally [3], and is the causative agent of trachoma (infectious blindness) in disadvantaged populations [4]. All members of the genus exhibit a unique biphasic developmental cycle where the non-replicating infectious elementary bodies (EBs) invade host cells and differentiate into replicating reticulate bodies (RBs) within a membrane-bound vacuole, escaping phagolysomal fusion [5]. *Chlamydia* actively modulates host cell processes to establish this intracellular niche, using secreted effectors and other proteins to facilitate invasion, internalisation and replication, while countering host defence strategies [6, 7]. At the end of the developmental cycle, RBs condense into EBs, which are released from the host cell by lysis or extrusion to initiate new infections [8].

Bacterial interactions with mammalian cells can induce dynamic transcriptional responses from the cell, either through bacterial modulation of host cell processes or from innate immune signalling cascades and other cellular responses [9–11]. In addition, effector proteins specifically targeting the nucleus (nucleomodulins) can influence cell physiology and directly interfere with transcriptional machinery including chromatin remodelling, DNA replication and repair [12]. Host cell epigenetic-mediated transcriptional regulatory changes, including histone modifications, DNA methylation, chromatin accessibility, RNA splicing, and non-coding RNA expression [13–15] may also be arbitrated by bacterial proteins and effectors. Consistent with host cell interactions with other bacterial pathogens, *C. trachomatis* infection alters host cell transcription over the course of its developmental cycle [16] and may also modulate the host cell epigenome. For example, NUE (<u>NU</u>clear <u>E</u>ffector), a *C. trachomatis* type III secreted effector with methyltransferase activity, enters the host nucleus and methylates eukaryotic histones H2B, H3 and H4 *in vitro* [17]. However, the ultimate gene targets of NUE activity or the affected host transcriptional networks are uncharacterised, as is the influence of chlamydial infection on the host cell epigenome in general.

To examine the impact of chlamydial infection on host cell chromatin dynamics, we applied FAIRE-Seq (Formaldehyde-Assisted Isolation of Regulatory Elements sequencing) [18] to *C. trachomatis*-infected HEp-2 epithelial cells and time-matched mock-infected cells, spanning the chlamydial developmental cycle (1, 12, 24 and 48 hours post infection). FAIRE protocols rely on the variable crosslinking efficiency of DNA to nucleosomes by formaldehyde, where nucleosome-bound DNA is more efficiently crosslinked. DNA fragments that are not crosslinked are subsequently enriched in the aqueous phase during phenol-chloroform extraction. These fragments represent regions of open chromatin, which in turn can be associated with regulatory factor binding sites. In FAIRE-Seq, libraries are generated from these enriched fragments, followed by sequencing and read mapping to a reference genome [18], allowing patterns of chromatin accessibility to be identified [19]. We identify infection-responsive changes in chromatin accessibility over the chlamydial developmental cycle, and identify several candidate host transcription factors that may be relevant to the cellular response to chlamydial infection.

## Methods

### Cell culture, infection and experimental design

HEp-2 cells (American Type Culture Collection, ATCC No. CCL-23) were grown as monolayers in 6 × 100mm TC dishes until 90% confluent. Monolayers were infected with *C. trachomatis* serovar E in SPG as previously described [20]. Additional monolayers were mock-infected with SPG only. The infection was allowed to proceed 48 hours prior to EB harvest, as previously described [20]. *C. trachomatis* EBs and mock-infected cell lysates were subsequently used to infect fresh HEp-2 monolayers. Fresh monolayers were infected with *C. trachomatis* serovar E in 3.5 mL SPG buffer for an MOI ~ 1 as previously described [20], using centrifugation to synchronize infections. Infections and subsequent culture were performed in the absence of cycloheximide or DEAE dextran. A matching number of HEp-2 monolayers were also mock-infected using uninfected cell lysates. Each treatment was incubated at 25°C for 2h and subsequently washed twice with SPG to remove dead or non-viable EBs. 10 mL fresh medium (DMEM + 10% FBS, 25μg/ml gentamycin, 1.25μg/ml Fungizone) was added and cell monolayers incubated at 37°C with 5% CO_2_. Three biological replicates of infected and mock-infected dishes per time were harvested post-infection by scraping and resuspending cells in 150μL sterile PBS. Resuspended cells were stored at −80°C.

We note that the experimental design used here cannot distinguish *Chlamydia*-mediated effects from infection-specific or non-specific host cell responses. Further experiments with inactivated *Chlamydia* or selected gene knock-outs or knock-downs will help to elucidate the extent of specific *Chlamydia*-mediated interference with the host cell epigenome. We also note that the use of *in vitro* immortalized HEp-2 epithelial cells means that, despite their utility and widespread use in chlamydial research, the full diversity of host cell responses that are likely to be found within *in vivo* infections will not be captured.

### FAIRE enrichment and sequencing

Formaldehyde-crosslinking of cells, sonication, DNA extraction of FAIRE-enriched fractions and Illumina library preparation was performed as previously described [18]. Libraries were sequenced on the Illumina 2500 platform at the Genome Resource Centre, Institute for Genome Sciences, University of Maryland School of Medicine.

### Bioinformatic analyses

Raw sequencing reads were trimmed and quality checked using Trimmomatic (0.36) [21] and FastQC (0.11.5) [22]. Trimmed reads were aligned to the human genome (GRCh 38.87) using Bowtie2 (2.3.2) [23] with additional parameters of ‘no mismatches’ and ‘–very-sensitive-local’. Duplicate reads were removed using Picard tools (2.10.4) [24]. Additional replicate quality control was performed using deepTools (2.5.3) [25] and in-house scripts.

Peak calling of open chromatin regions was performed using MACS2 (2.1.1) [26] in paired-end mode, with additional parameters of ‘–no-model –broad –q 0.05’ and MACS2 predicted extension sizes. All replicates were called separately, with significant peaks determined against the software-predicted background signal. Any peaks that fell within ENCODE blacklisted regions (regions exhibiting ultra-high signal artefacts) [27], or were located on non-standard chromosomes such as (ChrMT and ChrUn) were removed.

Consensus peak sets were created by combining significant peaks from the infected and mock-infected replicates for each time using Diffbind [28]. Peaks were removed if they appeared in less than two replicates. Reads were counted under each peak within each consensus peak set; the resulting read depths were normalised to their relative library sizes. Peaks with less than 3 mapped reads after normalisation were also removed. The resulting count matrices from each consensus peak set were used to look at the differences in chromatin accessibility between infected and mock-infected replicates at each time using the built in DESeq2 method of Diffbind (FDR < 0.05). This created a list of differential chromatin accessible regions, where patterns of open chromatin in either the mock-infected or infected conditions allowed corresponding patterns of closed chromatin to be identified in the matching condition. However, we note that, as FAIRE protocols are designed to enrich regions of open chromatin, there may be an inherent bias in favour of open chromatin.

Annotation of the set of differential chromatin accessible regions was performed with Homer (v4.9) [29] and separated into three main categories: Intragenic, Promoter and Intergenic. Intergenic: located >1kbp upstream of the transcriptional start site (TSS), or downstream from the transcription termination site (TTS); Promoter: located within 1kb upstream or 100bp downstream of the TSS (all promoter regions taken from RefSeq); and, Intragenic: annotated to a 3’UTR, 5’UTR, intron, exon, TTS, miRNA, ncRNA or a pseudogene. To identify enhancers, all intergenic regions were compared against experimentally validated enhancer regions from HeLa cells (S3 and S4) using Enhancer-atlas [30] and dbSuper [31].

Clusters of Open Regulatory Elements (COREs) were identified from the set of significant differential chromatin accessible regions using CREAM [32]. A window size of 0.5 and a peak range of 2:5 was initially set to separate COREs encompassing multiple genes from COREs overlapping individual genes. Subsequent filtering removed COREs with < 3 peaks and limited peak width to < 500,000 bp. Each CORE was visually inspected in the Integrative Genomics Viewer (IGV) to identify COREs that overlapped a single gene and to ensure all peaks had a fold-change of at least > 2 or < −2.

Motif analysis was performed with Homer [29]. Target sequences were regions with significant differential chromatin accessibility as identified by DESeq2, while the number of background sequences were randomly sampled regions throughout the human genome. Additional parameters included using a hypergeometric distribution, allowing for two mismatches and searching for motifs between 8-14 bp long. Motif enrichment was also performed with Homer [29], followed by filtering and assessment of human tissue specificity of the enriched transcription factors (TF) (p-value < 0.05, >5% of target sequences). For significant *de novo* TFs, motif matrices were compared against the Jaspar [33] and TomTom [34] databases, where enriched TFs were discarded unless the Homer annotation matched top hits in either database, and were also human-tissue specific.

Sequence data is available from the NCBI GEO archive GSE132448.

## Results and Discussion

### Chromatin accessibility landscapes of *Chlamydia*-infected and mock-infected cells

We applied FAIRE-Seq to *C. trachomatis* serovar E-infected and mock-infected human HEp-2 epithelial cells in triplicate at 1, 12, 24, and 48 hours post-infection (hpi). Following initial quality control measures, a single *C. trachomatis*-infected replicate was identified as an outlier and was removed from further analysis. The remaining replicates were mapped to the human genome (GRCh38), resulting in 52,584,839 mapped reads for mock-infected replicates and 98,802,927 mapped reads for *Chlamydia*-infected replicates (151,387,766 in total) (**Table 1**). Significant peaks, representing regions of open chromatin, were subsequently identified from these mapped reads. Each peak file was examined in IGV to ensure peaks were dispersed genome-wide without discernible chromosomal biases (**Additional File 1**). The total number of significant peaks from each replicate varied across the examined times and conditions, ranging between 1,759 and 17,450 peaks (**Figure 1A**).

**Table 1:**
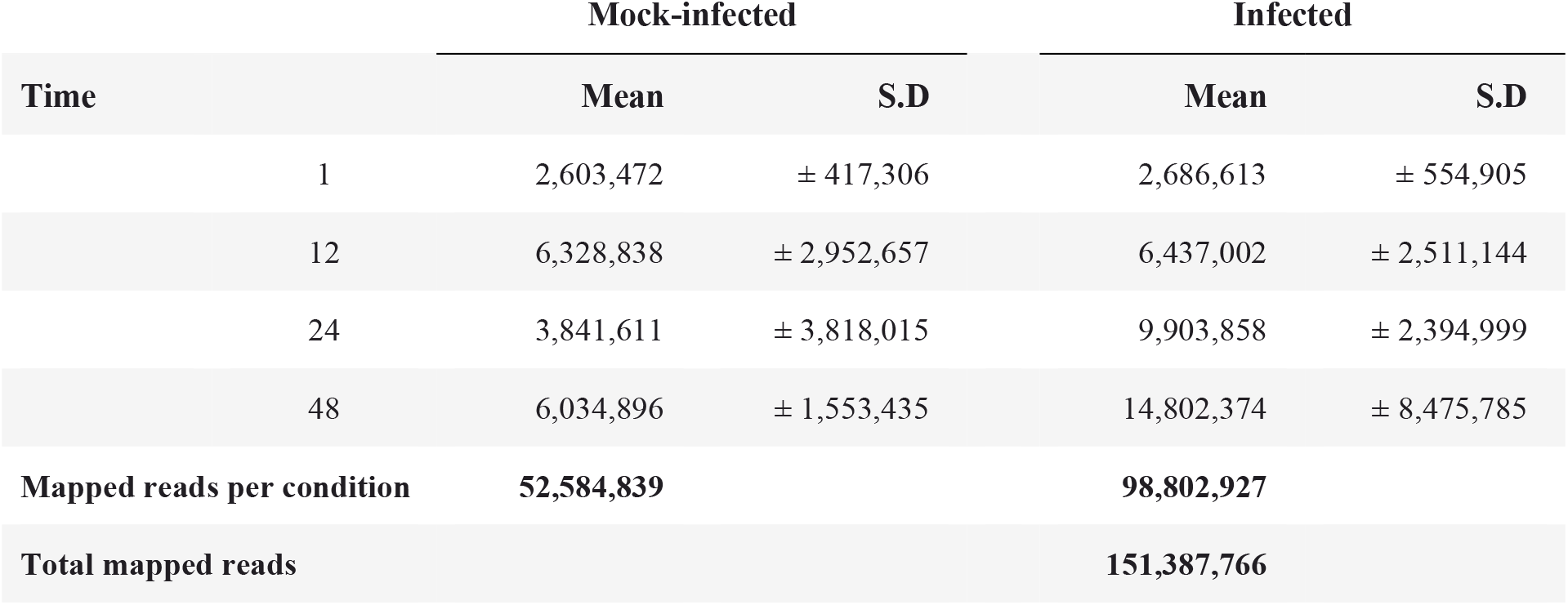
Summary of mapped reads, separated by time and condition

**Figure 1.**
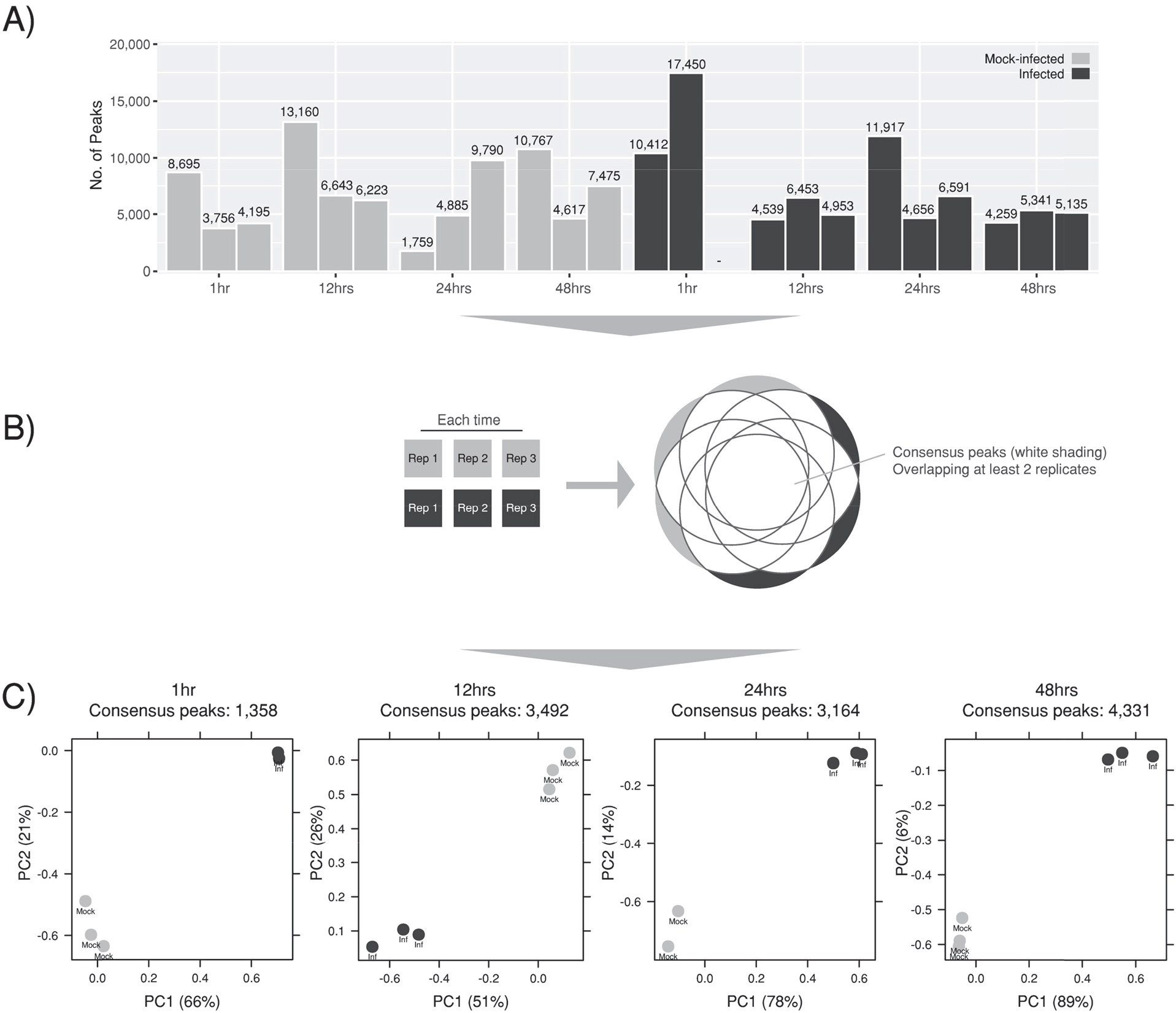
Identifying significant peaks and creating consensus peaksets. **A)** Significant peaks per replicate (p-value < 0.05). **B)** Consensus peaks were created for each time by combining significant peaks from *Chlamydia*-infected and mock-infected conditions, retaining peaks which appeared in > 2 replicates. **C)** PCA plots demonstrating tight clustering within each consensus peak set grouping infected and mock-infected replicates.

Diffbind [28] was used to group and filter peaks at each time post infection by removing regions with low coverage or any regions that were not represented across a consensus of replicates (**Figure 1B**). After normalisation for library size, principal component analysis (PCA) of the consensus peak sets (**Figure 1C**) led to the removal of one further outlier at 24 hours (mock-infected). The remaining peak sets exhibit tight clustering between mock-infected and infected conditions respectively at each time. Total consensus peak numbers increased across the chlamydial developmental cycle, independent of the total mapped reads over time.

### *C. trachomatis* infection is associated with temporal changes to chromatin accessibility in host cells

We identified genomic regions with significant differences in chromatin accessibility between infected and mock-infected conditions throughout the development cycle (FDR<0.05). The resulting set of differential chromatin accessible regions identifies both open and closed chromatin (relative to mock-infection). The total number of significant differentially accessible regions rose over the development cycle, with the number of regions increasing (3.6x) from 1 hpi (864) to 48 hpi (3,128) (**Figure 2A**). Open chromatin regions predominate at each time, (99% at 1hpi, 95% at 12 hpi, 97% at 24 hpi and 86% at 48 hpi) over closed chromatin regions, suggesting that host cell transcription and regulatory activity increases in response to infection.

**Figure 2.**
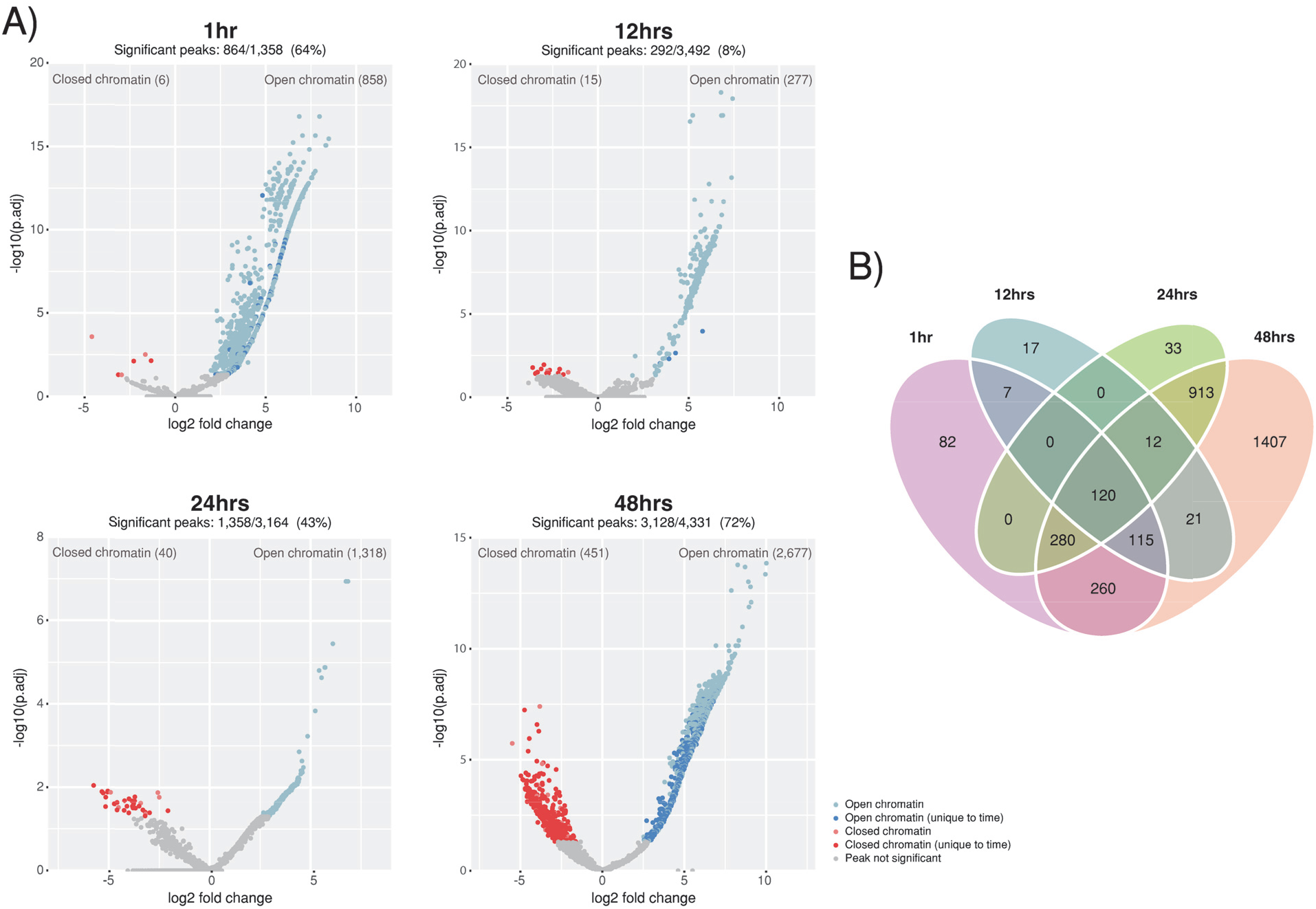
Changes in chromatin accessibility over the chlamydial developmental cycle. **A)** Volcano plots highlighting changes in chromatin accessibility between infected and mock-infected conditions. Regions of closed chromatin are represented as red dots, while open chromatin regions are blue dots. Peaks unique to a specific time have darker shading. Percentages above the plots show the proportion of consensus peaks with significant changes of chromatin accessibility between conditions (FDR < 0.05). **B)** Unique and conserved regions of differential chromatin accessibility across the developmental cycle.

At 12 hours, the number of significant differentially accessible regions was lower (8%), compared to the other times (64% at 1 hpi, 43% at 24 hpi and 72% at 48 hpi). The number of mapped reads was similar for all 12 hour replicates across conditions, and similar to other times, suggesting minimal bias from the variability of the underlying mapped reads (**Table 1**) and significant peaks (**Figure 1A**). In addition, each replicate had consistent peak coverage across the human genome (**Additional File 1**). Furthermore, 12 hour peak annotation is similar to other times (**Figure 3B-C**), and the distribution of peaks around the TSS (**Figure 3D**) are within promoter regions, as seen at 48 hours (**Figure 3D**). Thus, in the absence of any discernible bias, the lower number of significant differentially accessible regions at 12 hours may reflect a lower efficiency of formaldehyde crosslinking, or that this time in the course of chlamydial infection is relatively quiescent.

**Figure 3.**
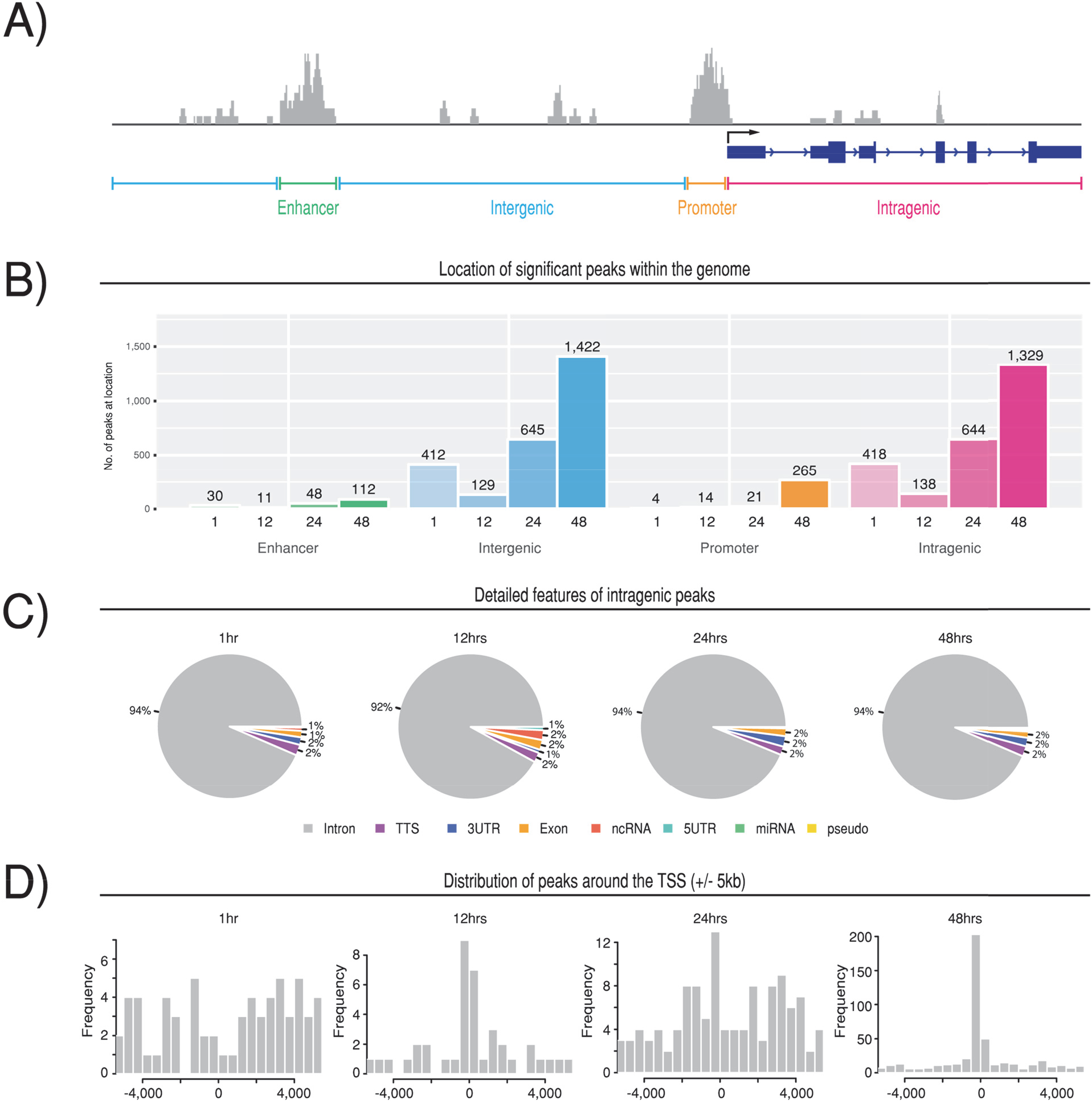
Annotation of significant peaks. **A)** Example illustration of annotating significant differential peaks to enhancer, promoter, intragenic or intergenic regions. **B)** Number of peaks per annotated category, separated by time. **C)** The intragenic peaks separated into eight detailed sub-categories. **D)** Distribution of all significant peaks and their proximity to the TSS of their associated genes (+/− 5KB).

120 differentially accessible chromatin regions are common at all examined times (**Figure 2B**), indicating a conserved response to chlamydial infection-associated events or general disruption of cellular homeostasis, irrespective of infection progression. Conversely, unique sets of differentially accessible regions are found at each time post-infection, highlighting the dynamism of the cellular response to infection over time, particularly at 48 hpi (**Figure 2B**). Most infection-associated differential chromatin accessible regions map to intergenic and intronic regions (**Figure 3B-C**, **Additional File 2**), consistent with other chromatin accessibility studies [35, 36], and the overall distribution of protein-coding genes within the human genome [37]. The distribution of differential chromatin-accessible regions around TSSs (+/− 5kb) at 12 and 48 hpi suggests that the majority of differential chromatin accessible regions are in proximity to TSSs. However, at 1 hpi there is no obvious distribution, while 24 hpi exhibits a bi-modal distribution (**Figure 3D**), suggesting that additional mechanisms, such as alternative splicing that may be contributing to the regulatory response to infection-associated events.

### Differential chromatin accessibility at promoters and enhancers identify infection-associated host regulatory activity

The proportion of all differentially accessible regions mapping to promoter regions is 4 (0.5%) at 1 hpi, 14 (4.8%) at 12 hpi, 21 (1.5%) at 24 hpi and 265 (8.5%) at 48 hpi (**Figure 4A**). Notably, 48 hpi exhibits a >10-fold increase in the number of significant regions compared to 24 hpi, with the majority of regions showing a reduction in chromatin accessibility, likely representing down-regulation of promoter-associated genes (**Figure 4A**). The large number of differentially accessible chromatin regions within promoters at 48 hours is a likely reflection of the diversity of events occurring at this late stage of the developmental cycle, including apoptosis, necrosis, lysis and cellular stress. Associated 48 hpi genes are linked with heat-shock stress (DNAJB1, DNAJB5, DNAJC21 and HSPA1B), cell defence (ILF2, MAP2K3 and STAT2), and cell stress/apoptosis (ATF3, PPM1B, GAS5, BAG1 and TMBIM6). ATP7A, which has a promoter exhibiting an increase in chromatin accessibility, is a key regulator of copper transport into phagosomes as part of a host cell response to intracellular infection [38, 39].

**Figure 4.**
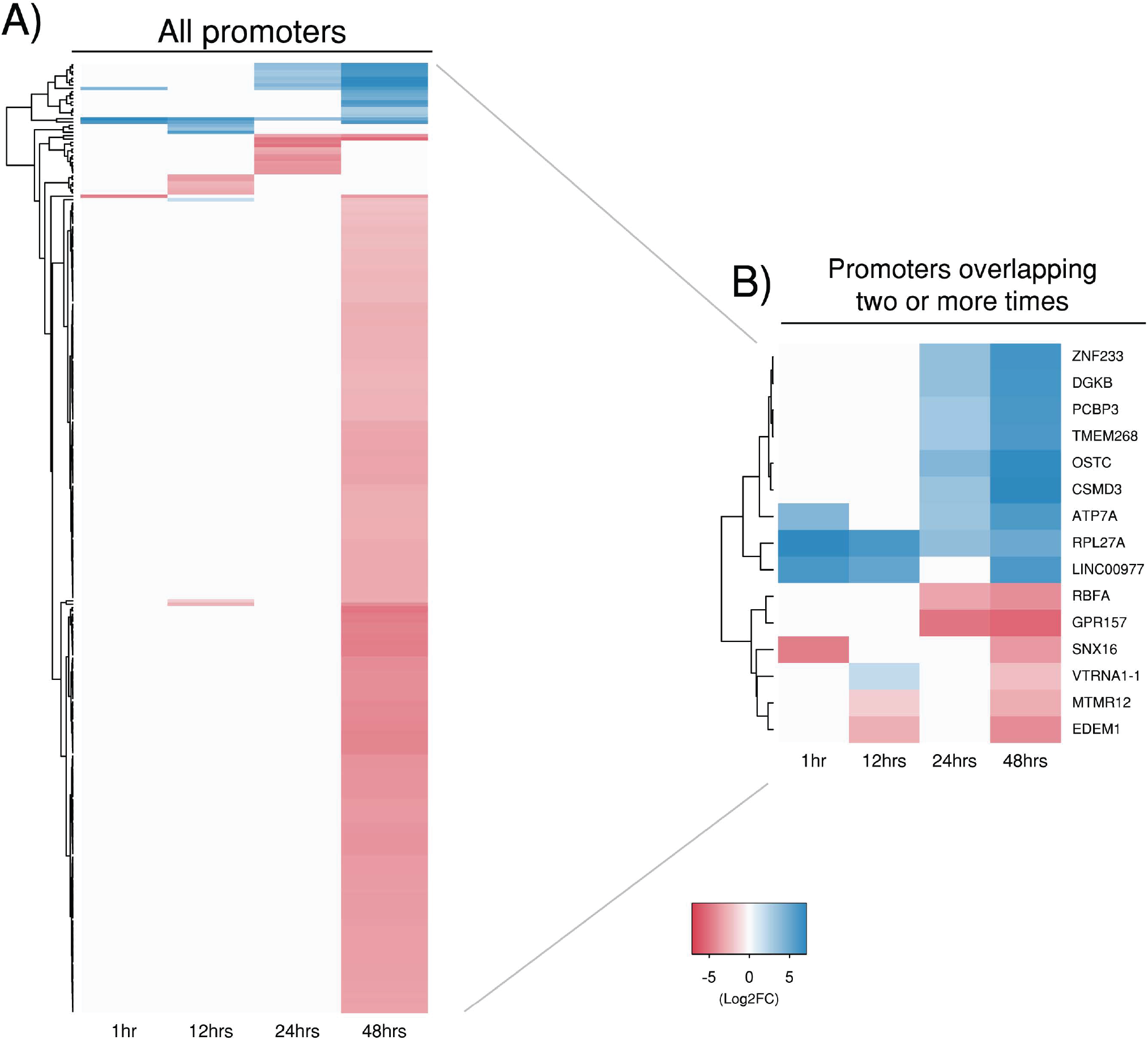
Differential chromatin accessibility within promoter regions. Heatmaps of significant differential peaks that were annotated to a promoter region. **A)** All promoter regions from each time post-infection. **B)** Promoters overlapping two or more times post-infection. Red and blue shading indicates fold-changes, while grey indicates no significant peaks.

Fifteen promoter-specific differentially accessible regions are found at two or more times. Two promoter regions are associated with genes encoding sorting nexin 16 (SNX16) and oligosaccharyltransferase complex subunit (OSTC) respectively (**Figure 4B**). The promoter region of OSTC exhibits increased chromatin accessibility at 24 and 48 hours; OSTC is linked to cellular stress responses [40]. Conversely, SNX16 shows a reduction in chromatin accessibility at both 1 and 48 hpi. Sorting nexins are a family of phosphatidylinositol binding proteins sharing a common PX domain that are involved in intracellular trafficking. Sorting nexins are a key component of retromer, a highly conserved protein complex that recycles host protein cargo from endosomes to plasma membranes or the Golgi [41]. Retromer is targeted by several intracellular pathogens, including *Chlamydia*, as a key strategy for intracellular survival [42]. The *C. trachomatis* effector protein, IncE, binds to sorting nexins 5 and 6, disrupting retromer-mediated host trafficking pathways [42] and potentially perturbing the endolysomal-mediated bacterial destruction capacity of the host cell [43]. However, SNX16 is a unique member of this family, containing a coiled-coil domain in addition to a PX domain, and is not associated with retromer [44]. SNX16 is instead associated with the recycling and trafficking of E-cadherin [44], which mediates cell-cell adhesion in epithelial cells, and is associated with a diversity of tissue specific processes, including fibrosis and epithelial-mesenchymal transition (EMT) [45]. Separately, *C. trachomatis* infection has been shown to downregulate E-cadherin expression via increased promotor methylation, potentially contributing to EMT-like changes [46]. Thus, downregulation of SNX16, as inferred by the observed reduction in promotor-associated chromatin accessibility may contribute to chlamydial fibrotic scarring outcomes. In other bacterial pathogens, modulation of E-cadherin is a known virulence mechanism where it is degraded by proteases, such as HtrA, disrupting tight and adherens junctions to facilitate invasion through the epithelial barrier [47, 48]. Although chlamydial HtrA has been detected outside the inclusion and in exported blebs [49], E-cadherin has not yet been identified as a chlamydial HtrA target. Nevertheless, HtrA has been shown to be critical for *in vivo* chlamydial infections, indicating that this functionality may be revealed in the future [50].

Changes in chromatin accessibility of regions overlapping tissue-specific enhancers from experimentally validated databases were examined, identifying 211 enhancers and seven “super-enhancers” (**Figure 5A**). All super-enhancers exhibited an increase in chromatin accessibility, and were associated with genes mediating cell growth (KLF5), cell structure and signalling (FLNB, PTP4A2 and MSN), and innate immunity (IER3) (**Additional File 3**). Infection-responsive chromatin accessible regions occurring at three or more times over the chlamydial developmental cycle (all exhibiting an increase in chromatin accessibility) identified known enhancers that influence DNA/RNA-polymerase activity (AFF1, POLR2M, TCEB1, CHMP4C and POLL), including elongation factors, chromatin remodelling and DNA repair (**Figure 5B**). The manipulation of these genes and underlying functions are suggestive of nucleomodulin activity, which are a class of bacterial effectors that directly target the host cell nucleus to manipulate host defences and machinery [12]. One example of a *C. trachomatis* specific nucleomodulin is NUE, which is directed to the nucleus and performs methyltransferase activity [17]. However, as noted above, our experimental design does not distinguish *Chlamydia*-mediated effects from infection-specific or non-specific host cell responses.

**Figure 5.**
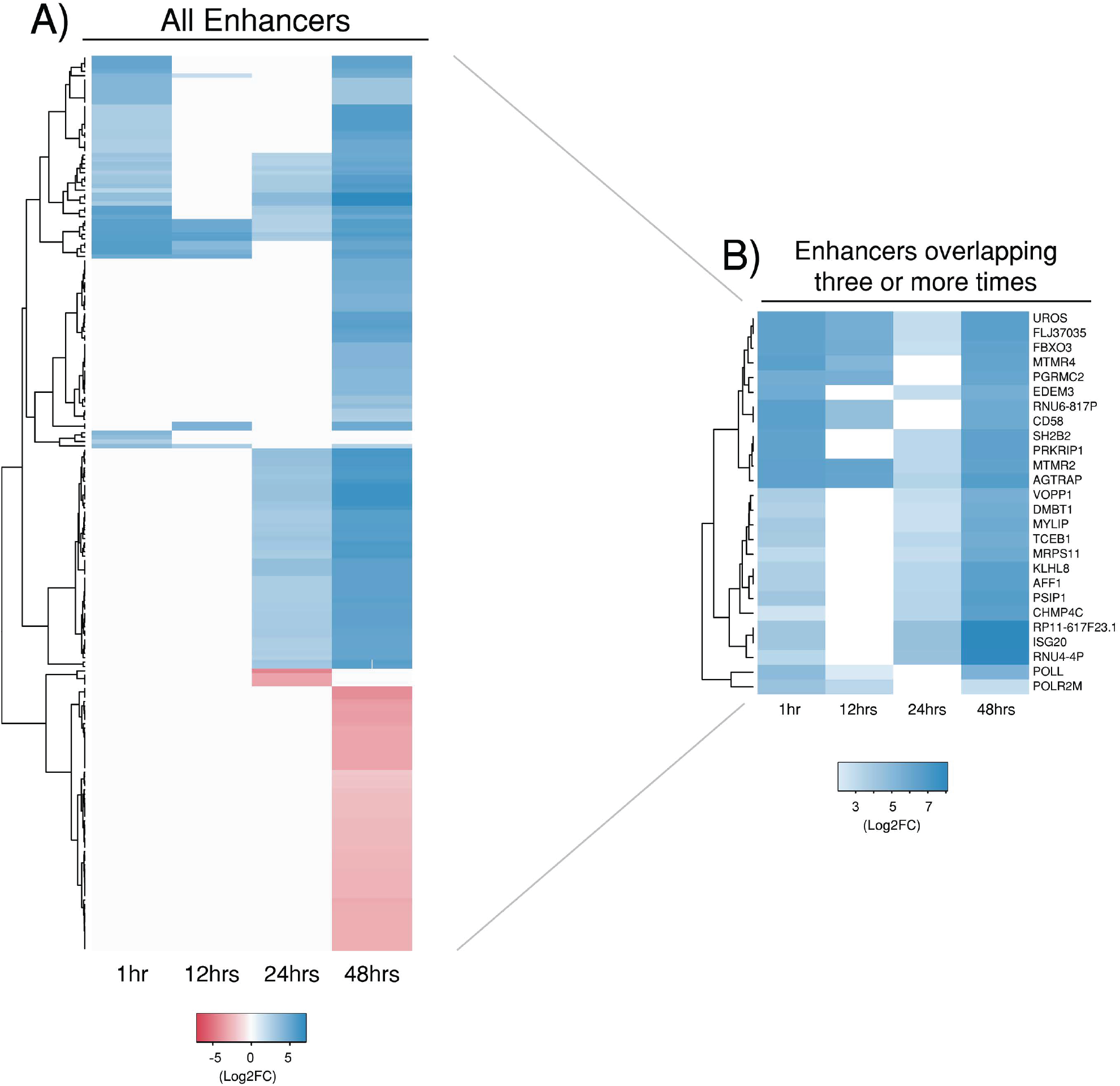
Differential chromatin accessibility within enhancer regions. Significant differential peaks annotated as intergenic were compared against experimentally validated tissue-specific enhancers. **A)** All enhancer regions across each time. Seven super enhancers were identified and are denoted with a star (*). **B)** Enhancers overlapping three or more times. Red and blue shading indicate fold-changes, while grey indicates that no significant peaks were associated with that enhancer. Some enhancers contain more than one peak, explaining why there are multiple fold-changes at some times.

In addition, three enhancer-linked genes that recur three or more times over the developmental cycle and show an increase in chromatin accessibility, are involved in ubiquitination and protein quality control (KLHL8, FBXO3 and EDEM3). The eukaryotic ubiquitination modification marks proteins for degradation and regulates cell signalling of a variety of cellular processes, including innate immunity and vesicle trafficking [51]. The deposition of ubiquitin onto intracellular pathogens is a conserved mechanism found in a diverse range of hosts [52]. In *Chlamydia*, host cell ubiquitin systems can mark chlamydial inclusions for subsequent destruction [53] and there is emerging evidence that various *Chlamydia* species, using secreted effectors and other proteins, are able to subvert or avoid these host ubiquitination marks for intracellular survival [53, 54]. Our observation of increased chromatin accessibility of enhancer elements linked to ubiquitination genes, putatively augmenting expression of these genes, further highlights the complex role of ubiquitination in chlamydial infection.

### Conserved and time-specific host responses to infection over the chlamydial developmental cycle

Differential chromatin accessible regions that are present at all four times during infection demonstrate a conserved host cell response to chlamydial infection (**Figure 2B**). Time-specific differential chromatin accessibility is also evident over the chlamydial developmental cycle (**Figure 2B**). To investigate the conserved host cell response, we focused upon 63 of the 120 differential chromatin accessible regions (intragenic, promoter or enhancer regions) identified above, excluding the likely ambiguous intergenic regions (**Figure 6A**). 56 were within intronic regions, one within a 3’UTR (FECH), a promoter (RPL27A), and five within enhancer regions (MTMR2, FLJ37035, UROS, FBXO3 and AGTRAP). Only 4 of these 63 significant differentially accessible regions show a decrease in overall chromatin accessibility. However, these same regions also exhibit increased chromatin accessibility at different intragenic locations at 48 hpi, further highlighting the potential for infection-related alternative splicing mechanisms (**Figure 6A**). The remaining conserved differentially accessible regions were associated with genes involved in infection-relevant cellular processes, including C8A as part of the complement cascade, and lipase activity from LIPI that is essential for chlamydial replication [55]; while multiple genes (HDAC2, HNRNPUL1, NCOA7 and YAP1) are known transcriptional regulators. We also examined any differential chromatin accessible regions that appeared across three times. This identified further effects of infection on the complement cascade. Key components of the membrane attack complex (MAC) and complement activation pathways exhibit increased differential chromatin accessibility (C8B at 1, 12 and 24 hours and CFHR5 at 24 and 48 hours). Conversely, C6 exhibits decreased chromatin accessibility at 48 hours.

**Figure 6.**
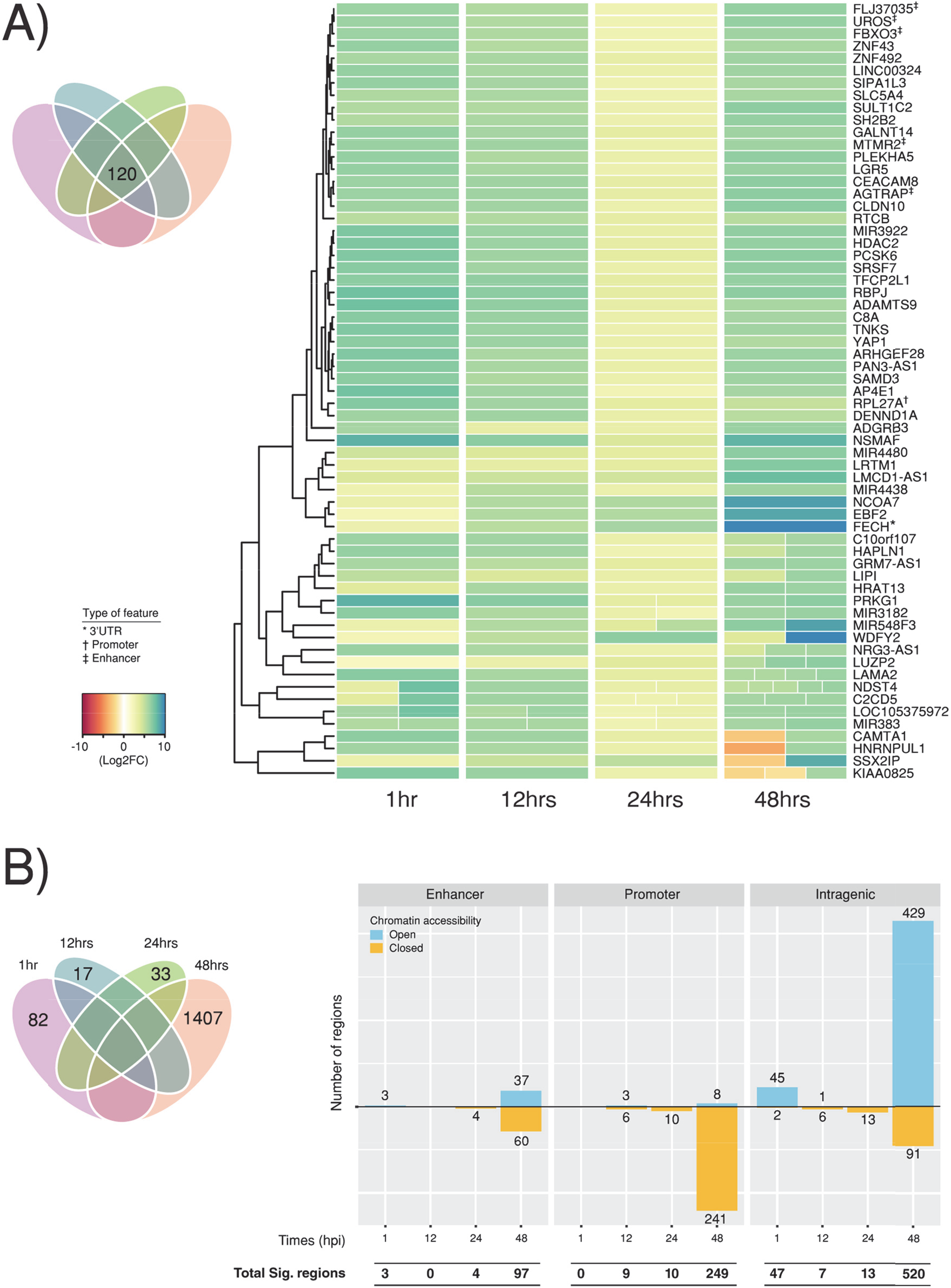
Conserved host cell response to infection. A) 120 Differentially accessible regions found in all four times were extracted, representing a conserved host cell response to infection. Intergenic regions were removed due to the ambiguity of annotating to the closest feature. If a gene contained more than one peak within a specific time, the different fold changes are split out evenly within the column at that time. B) Venn diagram highlighting the number of time-specific differential regions. Intergenic regions were also removed for the same reasons, with the remaining enhancers, promoters and intragenic regions separated by their chromatin accessibility.

We identified unique differentially accessible regions across the chlamydial developmental cycle (**Figure 6B**). At 1, 12 and 24 hpi, there are a relatively small number of significant differential chromatin accessible regions. In contrast, 48 hpi exhibits over 1,400 regions, further reflecting the diverse processes associated with the end of the *in vitro* developmental cycle as indicated previously. As with the conserved differential regions above, we focused on differential chromatin accessibility within promoters, enhancers and intragenic regions (50 at 1 hpi, 17 at 12 hpi, 27 at 24 hpi and 866 at 48 hpi) (**Figure 6B**, **Additional File 4)**.

At 1 hpi, increased chromatin accessibility was associated with a variety of genes involved in the regulation of host cell defences (CD44, IFNAR1, LGALS8, STAT1, SLA2 and DDAH1), transcription and translation (ZNF461, ZNF800, PHF2, PABPC4L, RPS13 and SIN3A), the cell cycle (NIPBL, CEP57L1 and CMTM4) and BCL2L14 (Apoptosis facilitator Bcl-2-like protein 14) a member of the Bcl-2 Family of proteins that are linked to apoptosis [56] (**Figure 7A**). At 12 hours, four ncRNAs were identified (RPPH1, RN7SK, RN7SL2 and RMRP) that are involved in RNA processing, signalling and transcriptional regulation [57-60]. The remaining genes at 12 hours exhibited decreased chromatin accessibility, encompassing the cell cycle and DNA replication (SDCCAG8 and ORC2), and ubiquitination (PJA2 and FBXO46) (**Figure 7B**). At 24 hours, all genes were associated with decreased chromatin accessibility and were grouped into four sub-categories: cell cycle (WAPL, SMARCB1 and CDC20), energy production (HK1, ACO1 and SLC25A13), metabolism (ARSA, EXTL3 and SLC27A2), and transcription (AP5Z1 and ELP3) (**Figure 7C**).

**Figure 7.**
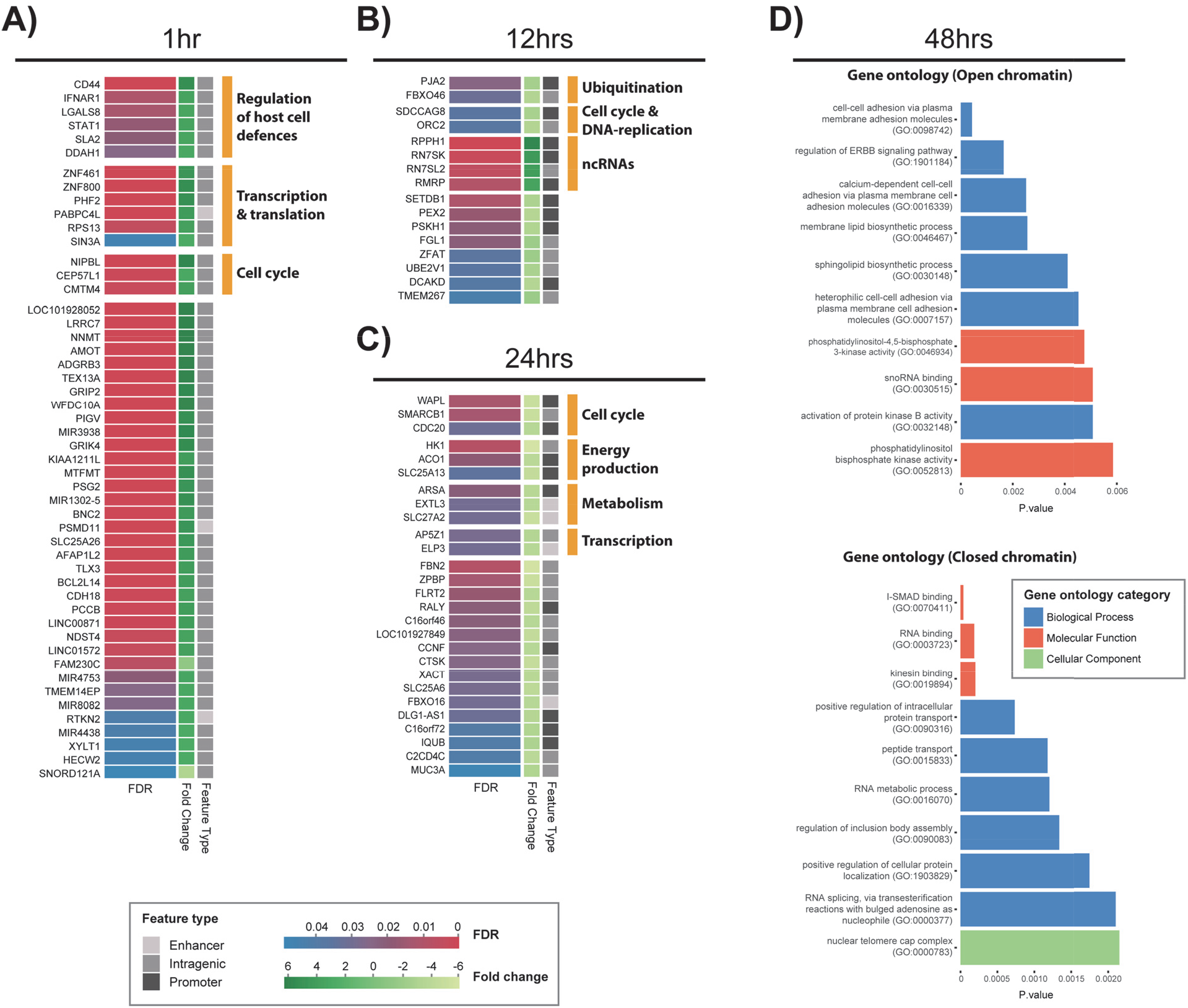
Enrichment of time-specific differential chromatin regions. Annotated time-specific differential chromatin regions associated with 1 hour **A)**, 12 hours **B)** and 24 hours **C)**. Where genes have been grouped into annotated categories, multiple underlying sources were used for verification. **D)** At 48 hours, a substancial increase in genes allowed Gene Ontology (GO) enrichment. All three GO categories were enriched, with the top ten p-values across the categories displayed.

### Increased changes to differential chromatin accessibility at the end of the developmental cycle

The large number of genes associated with differential chromatin accessibility at 48 hours permitted Gene Ontology enrichment to be performed, with the underlying genes distinguished by increased chromatin and decreased chromatin accessibility (**Figure 7D**). Significantly enriched ontologies associated with regions of increased chromatin accessibility include the ErbB signalling pathway *(GO:1901184)*, which is linked to a wide range of cellular functions including growth, proliferation and apoptosis. ErbB transmembrane receptors are also often exploited by bacterial pathogens for host cell invasion [61]. Notably, epidermal growth factor receptor (EGFR), a member of the ErbB family, is the target receptor for *C. pneumoniae* Pmp21 as an EGFR-dependent mechanism of host cell entry [62]. The *C. trachomatis* Pmp21 ortholog, PmpD, also has adhesin-like functions [63], however the host ligands are unknown. Nevertheless, EGFR inhibition results in small, immature *C. trachomatis* inclusions, with calcium mobilisation and F-actin assembly disrupted [64], indicating the functional importance of EGFR and the ErbB signaling pathway for *C. trachomatis* attachment and development.

Three enriched biological processes share the term ‘*cell-cell adhesion via plasma membrane adhesion molecules*’ (*GO:0098742, GO:0016339 and GO:0007157*). Several genes common to these categories with infection-responsive differential chromatin accessibility are associated with cadherins (CDH4, CDH12, CDH17, CDH20, FAT4 and PTPRD). Disruption of cadherin function has been described in *C. trachomatis* infection, and is linked to the alteration of adherens junctions and the induction of epithelial-mesenchymal transition (EMT) events that may underlie chlamydial fibrotic outcomes [46, 65]. Altered chromatin accessibility for other cadherin-relevant loci over the chlamydial developmental cycle is apparent in this data, including SNX16 (see above) and CDH18 (see below), suggesting that alteration or disruption of cadherin regulation is a key feature of chlamydial infection. Two enriched lipid-based biological processes, ‘*Sphingolipid biosynthesis (GO:0030148)*, and ‘*Membrane lipid biosynthetic process* (*GO:0030148*) are also associated with regions of open chromatin. *Chlamydia* scavenges a range of host-cell-derived metabolites for intracellular growth and survival, particularly lipids [66, 67].

Significantly enriched ontologies associated with regions of decreased chromatin accessibility include the ‘*I-Smad (inhibition of Smad) binding*, *(GO:0070411)’*. I-Smads (Inhibitory-Smads) are one of three sub-types of Smads that inhibit intracellular signalling of TGF-β by various mechanisms including receptor-mediated inhibition [68]. In addition, Smad2 contains two closed chromatin accessibility regions at an enhancer and intragenically respectively. Smad2 is part of the R-Smad sub-family that regulates TGF-β signalling directly [69, 70]. TGF-β has been hypothesised to be a central component of dysregulated fibrotic processes in *Chlamydia*-infected cells, provoking runaway positive feedback loops that generate excessive ECM deposition and proteolysis, potentially leading to inflammation and scarring [16]. We also identify down-regulation of the ontology *‘Kinesin binding (GO:0019894)*. Kinesins belong to a class of motor proteins that move along microtubule filaments (from the centre of the cell outwards) supporting cell functions including transport and cell division [71]. *C. trachomatis* expresses an effector protein (CEP170) that recruits host microtubules into the vicinity of the mature inclusion, enabling microtubule-dependent traffic to be re-directed to the inclusion [72].

### Clusters of Open Regulatory Elements

Clusters of Open Regulatory Elements (COREs) are multiple areas of open chromatin in close proximity to each other, which may represent regions of coordinated chromatin accessibility linked to multiple regulatory elements [73]. We focused on differential chromatin regions spanning less than 500k bp that contain a cluster of at least three regions (**Figure 8A**). This identified 18 COREs across three times post-infection consisting of regions with the same fold-change direction and overlapping a single gene (**Figure 8B**). A CORE is apparent at 1 hpi in proximity to laminin (LAMA2). Laminins are a component of the extracellular matrix and basement membranes that influence cell differentiation, migration, and adhesion. As noted above, dysregulation of ECM moieties has been hypothesised to be a key mechanism of chlamydial scarring, in which immune-mediated positive feedback loops are induced on infection as part of an early, aberrant wound response to chlamydial infection, creating inflammatory accumulations of ECM constituents [16]. Combined with the observed chromatin accessibility changes to several cadherin and cadherin-associated genes and TGF-β-mediated Smad signalling in this work, a CORE within the laminin gene provides further support for the key role of dysregulated ECM in chlamydial disease outcomes.

**Figure 8.**
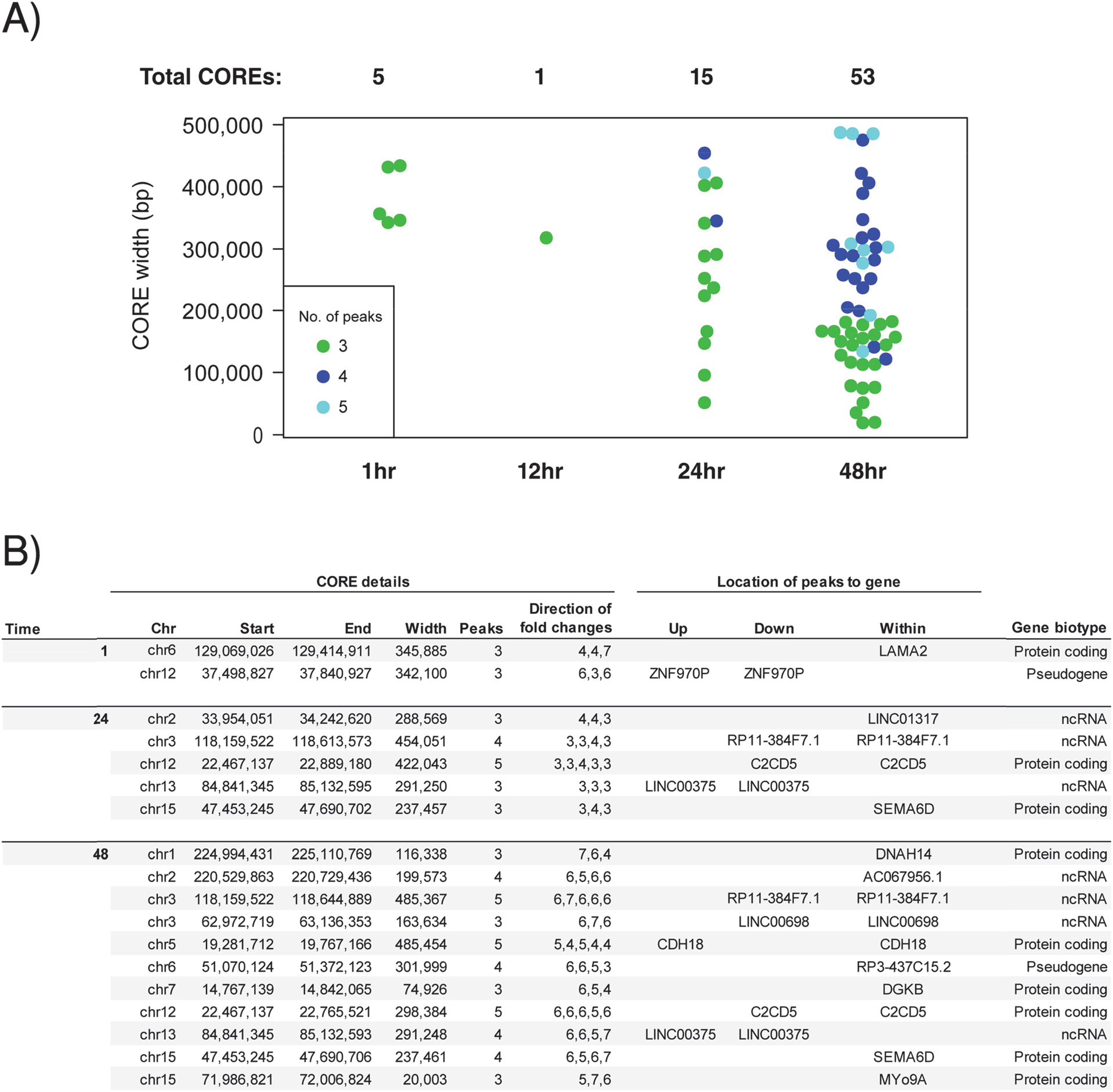
COREs (Clusters of Open Regulatory Elements) **A)** Number of COREs at each time post-infection using significant differential peaks, separated by width and the number of peaks within each CORE. COREs have a maximum width of 500,000 bp and > 3 peaks. **B)** 18 significant COREs were identified across three times post-infection. For each CORE, the genomic location, associated number of peaks, where they fall within proximity to a genomic feature, fold-changes, and genetic biotype are shown.

At 48 hours, eleven COREs were identified, overlapping six protein-coding and five non-coding genes. Two of these genes (DNAH14 and MYo9A) belong to broad families of cytoskeletal motor proteins (dyneins and myosins), with relevance to chlamydial infection. Some members of the myosin family may be involved with chlamydial extrusion through a breakdown of the actin cytoskeleton followed by the release of EB’s at the end of the lifecycle [74]. However, MYo9A itself has not been previously linked to chlamydial infection. Similarly, dynein-based motor proteins have been shown to move the chlamydial inclusion via the internal microtubule network to the MTOC (Microtubule-Organizing Centre); the close proximity to the MTOC is thought to facilitate the transfer of host vesicular cargo to the chlamydial inclusion [75]. However, DNAH14 is an axonemal dynein that causes sliding of microtubules in the axonemes of cilia and flagella, and is typically only expressed in cells with those structures [76]; it is not clear what role it would have in chlamydial infection. A third CORE overlaps DGKB, a diacylglycerol kinase that metabolises 1,2,diacylglycerol (DAG) to produce phosphatidic acid (PA), a key precursor in the biosynthesis of triacylglycerols and phospholipids, and a major signalling molecule [77]. *Chlamydia* obtains and redirects host-derived lipids through multiple pathways [78], and as further identified in this CORE and enriched gene ontologies (above).

### Identification of transcription factor motifs

Putative transcription factor (TFs) motifs were identified from all significant differential chromatin accessible regions at each time post-infection (**Additional File 5**). Ten significant TF motifs were identified, spanning the developmental cycle (**Table 2**). IRF3 (Interferon Regulatory Factor) motifs are enriched at 1 hpi; IRF3 is a key transcriptional regulator of type I interferon (IFN)-dependent innate immune responses and is induced by chlamydial infection. The type I IFN response to chlamydial infection can induce cell death or enhance the susceptibility of cells to pro-death stimuli [79], but may also be actively dampened by *Chlamydia* [80, 81]. Specificity Protein 1 (Sp1) is a zinc-finger TF that binds to a wide range of promoters with GC-rich motifs. Sp1 may activate or repress transcription in a variety of cellular processes that include responses to physiological and pathological stimuli, cell differentiation, growth, apoptosis, immune responses, response to DNA damage and chromatin remodelling [82, 83].

**Table 2:**
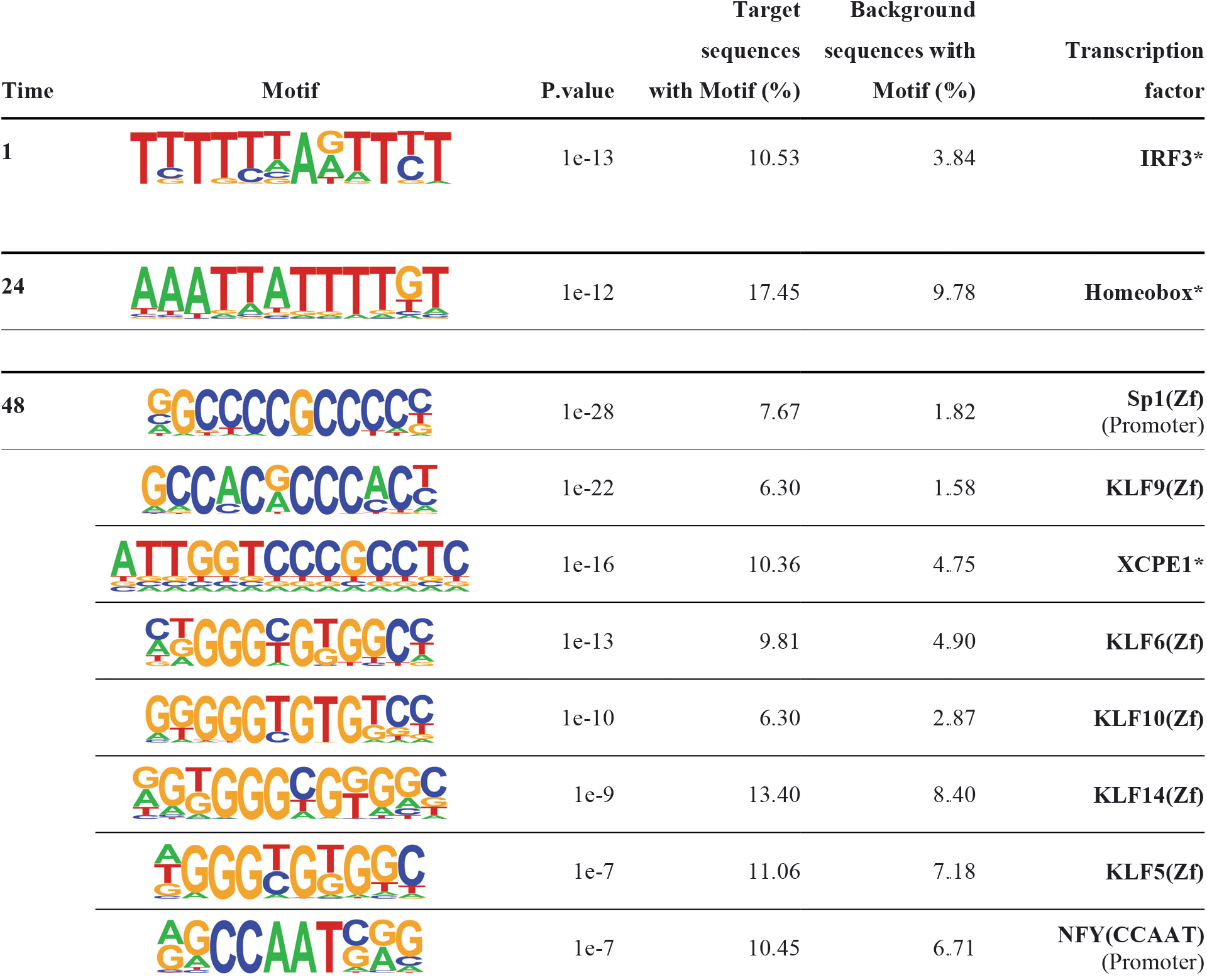
Motifs and enriched transcription factors. Target sequences are significant differential peaks and background sequences are randomly selected throughout the genome to determine significance. A star (*) denotes a de-novo motif where various sources were used to annotate the corresponding transcription factor.

| Time | Motif | P.value | Target sequences with Motif (%) | Background sequences with Motif (%) | Transcription factor |
| --- | --- | --- | --- | --- | --- |
| 1 |  | 1e-13 | 10.53 | 3.84 | IRF3* |
| 24 |  | 1e-12 | 17.45 | 9.78 | Homeobox* |
| 48 |  | 1e-28 | 7.67 | 1.82 | Sp1(Zf)<br>(Promoter) |
|  |  | 1e-22 | 6.30 | 1.58 | KLF9(Zf) |
|  |  | 1e-16 | 10.36 | 4.75 | XCPE1* |
|  |  | 1e-13 | 9.81 | 4.90 | KLF6(Zf) |
|  |  | 1e-10 | 6.30 | 2.87 | KLF10(Zf) |
|  |  | 1e-9 | 13.40 | 8.40 | KLF14(Zf) |
|  |  | 1e-7 | 11.06 | 7.18 | KLF5(Zf) |
|  |  | 1e-7 | 10.45 | 6.71 | NFY(CCAAT)<br>(Promoter) |

The majority of TF motifs enriched at 48 hours correspond to Krüppel-like-factors (KLFs). KLFs are zinc-finger TFs in the same family as Sp1, which is also enriched at 48 hours. The members of this large family orchestrate a range of paracrine and autocrine regulatory circuits and are ubiquitously expressed in reproductive tissues [84]. Dysregulation of KLFs and their dynamic transcriptional networks is associated with a variety of uterine pathologies [85]. We find motif enrichment for five distinct KLFs (KLF5, KLF6, KLF9, KLF10 and KLF14) at 48 hours, in addition to further KLFs at 12 and 48 hours (KLF 3 and KLF 4) without the initial filtering steps (**Additional File 5**). KLF5 is a transcriptional activator found in various epithelial tissues and is linked to regulation of inflammatory signalling, cell proliferation, survival and differentiation [86]. KLF6 is also a transcriptional activator ubiquitously expressed across a range of tissues and plays a crucial role in regulating genes involved with tissue development, differentiation, cell cycle control, and proliferation [87]. Target genes include collagen α1, keratin 4, TGFβ type I and II receptors, and others [88]. KLF9, 10 and 14 act as transcriptional repressors and are ubiquitously expressed across a range of tissues [89]. KLF9 is a tumour suppressor [90] and regulates inflammation, while KLF10 has a major role in TGF-β-linked inhibition of cell proliferation, inflammation and initiating apoptosis [91]. KLF14 represses TGF-βRII activity in inflammation [92], regulates lipoprotein metabolism [93], and is induced upon activation of naïve CD4+ T cells [94].

Histone deacetylases (HDACs) modify the core histones of the nucleosome, providing an important function in transcriptional regulation [95], and many bacterial pathogens subvert HDACs to suppress host defences [15]. KLF9, 10 and 14 share the co-factor Sin3A (SIN3 Transcription Regulator Family Member A) [60], which is also a core component of the chromatin-modifying complex mediating transcriptional repression [66]. The Sin3a/HDAC complex is made up of two histone deacetylases HDAC1 and HDAC2. HDAC2 has increased chromatin accessibility at all four time points, and HDAC9 has increased chromatin accessibility at 1, 24 and 48 hours, further supporting the potential for histone modifications to be a component of the host cell response to chlamydial infection, or to be targets of chlamydial effectors [17].

## Conclusions

We describe comprehensive changes to chromatin accessibility upon chlamydial infection in epithelial cells *in vitro*. We identify both conserved and time-specific infection-responsive changes to a variety of features and regulatory elements over the course of the chlamydial developmental cycle that may shape the host cell response to infection, including promotors, enhancers, COREs, and transcription factor motifs. Some of these changes are associated with genomic features and genes known to be relevant to chlamydial infection, including innate immunity and complement, acquisition of host cell lipids and nutrients, intracellular signalling, cell-cell adhesion, metabolism and apoptosis. Host cell chromatin accessibility changes are evident over the entire chlamydial developmental cycle, with a large proportion of all chromatin accessibility changes at 48 hours post infection. This likely reflects the confluence of late stages of developmental cycle events, however significant changes to chromatin accessibility are readily apparent as early as 1 hour post infection. We find altered chromatin accessibility in several gene regions, ontologies and TF motifs associated with ECM moieties, particularly cadherins and their interconnected regulatory pathways, laminin, and Smad signalling. Disruption of the ECM is thought to be a central component of dysregulated fibrotic processes that may underpin the inflammatory scarring outcomes of chlamydial infection [16], and our data further highlights a central role of the ECM in epithelial cell responses to infection. We also identify factors that have not been previously described in the context of chlamydial infection, notably the enrichment of the KLF family of transcription factor motifs within differential chromatin accessible regions in the latter stages of infection. Dysregulation of the biologically complex KLFs and their transcriptional networks is linked to several reproductive tract pathologies in both men and women [85], thus our discovery of enriched KLF binding motifs in response to chlamydial infection is compelling, given the scale and burden of chlamydial reproductive tract disease globally [3].

In summary, this is the first genome-scale analysis of the impact of chlamydial infection on the human epithelial cell epigenome, encompassing the chlamydial developmental cycle at early, mid and late times. This has yielded a novel perspective of the complex host epithelial cell response to infection, and will inform further studies of transcriptional regulation and epigenomic regulatory elements in *Chlamydia*-infected human cells and tissues. Examination of the multifaceted human epigenome, and its potential subversion by *Chlamydia*, using *in vivo* mouse models of infection and *ex vivo* human reproductive tract tissues, will continue to shed light on how the host cell response contributes to infection outcomes.

## Supporting information

Additional File 1.docx

Additional File 3.docx

## Acknowledgements

Sequencing was performed at the Genome Resource Center, Institute for Genome Sciences, University of Maryland School of Medicine and funded by the Genome Sequencing Center for Infectious Diseases (NIAID HHSN272200900009C). This research was supported by UTS Faculty of Science Startup funding to GM. Data was analysed on the ARCLab high-performance computing cluster at UTS, with files hosted using the Spaceshuttle facility at Intersect Australia.

## Additional Files

**Additional File 1.docx Genome coverage plots**

Significant peaks from each replicate as determined by MACS2. Screenshots are from IGV (Integrative Genomics Viewer) showing that all replicates contain significant peaks genome-wide (human genome) without any visual chromosomal bias.

**Additional File 2.xlsx Annotation of all significant peaks**

Annotation of all the significant peaks, with tabs separating genomic features and fold-change regulation.

**Additional File 3.docx Enrichment of Super-enhancer genes**

Super enhancer-linked genes separated by time and biological activity.

**Additional File 4.xlsx Time specific regions**

The list of time-specific differential chromatin accessible regions. It should be noted that some genes in these lists are repeated at each time due to multiple peaks occurring at an annotated interval, that enhancers can affect more than one gene, and single genes can be affected by more than one enhancer.

**Additional File 5.xlsx Complete list of motifs and transcription factors**

The complete list of significant motifs and enriched transcription factors.

## References

1. Schachter J, Storz J, Tarizzo ML, Bögel K: Chlamydiae as agents of human and animal diseases. Bulletin of the World Health Organization 1973, 49(5):443–449.

2. Reyburn H: WHO Guidelines for the Treatment of Chlamydia trachomatis. WHO 2016, 340(may28 1):c2637–c2637.

3. Menon S, Timms P, Allan JA, Alexander K, Rombauts L, Horner P, Keltz M, Hocking J, Huston WM: Human and Pathogen Factors Associated with Chlamydia trachomatis-Related Infertility in Women. Clinical microbiology reviews 2015, 28(4):969–985.

4. Burton MJ: Trachoma: an overview. British Medical Bulletin 2007, 84(1):99–116.

5. Fields KA, Hackstadt T: The Chlamydial Inclusion: Escape from the Endocytic Pathway. Annual Review of Cell and Developmental Biology 2002, 18(1):221–245.

6. Dautry-Varsat A, Balana ME, Wyplosz B: Chlamydia--host cell interactions: recent advances on bacterial entry and intracellular development. Traffic (Copenhagen, Denmark) 2004, 5(8):561–570.

7. Betts-Hampikian HJ, Fields KA: The Chlamydial Type III Secretion Mechanism: Revealing Cracks in a Tough Nut. Frontiers in microbiology 2010, 1:114.

8. Hybiske K, Stephens RS: Mechanisms of host cell exit by the intracellular bacterium Chlamydia. Proceedings of the National Academy of Sciences of the United States of America 2007, 104(27):11430–11435.

9. Brunham RC, Rey-Ladino J: Immunology of *Chlamydia* infection: implications for a *Chlamydia trachomatis* vaccine. Nat Rev Immunol 2005, 5(2):149–161.

10. Alonso A, Garcia-del Portillo F: Hijacking of eukaryotic functions by intracellular bacterial pathogens. International microbiology : the official journal of the Spanish Society for Microbiology 2004, 7(3):181–191.

11. Ribet D, Cossart P: How bacterial pathogens colonize their hosts and invade deeper tissues. Microbes and Infection 2015, 17(3):173–183.

12. Bierne H, Cossart P: When bacteria target the nucleus: the emerging family of nucleomodulins. Cellular microbiology 2012, 14(5):622–633.

13. Bierne H, Hamon M, Cossart P: Epigenetics and bacterial infections. Cold Spring Harbor perspectives in medicine 2012, 2(12):a010272.

14. Hamon MA, Cossart P: Histone modifications and chromatin remodeling during bacterial infections. Cell Host Microbe 2008, 4(2):100–109.

15. Grabiec AM, Potempa J: Epigenetic regulation in bacterial infections: targeting histone deacetylases. Critical Reviews in Microbiology 2018, 44(3):336–350.

16. Humphrys MS, Creasy T, Sun Y, Shetty AC, Chibucos MC, Drabek EF, Fraser CM, Farooq U, Sengamalay N, Ott S et al: Simultaneous Transcriptional Profiling of Bacteria and Their Host Cells. PLOS ONE 2013, 8(12):e80597.

17. Pennini ME, Perrinet S, Dautry-Varsat A, Subtil A: Histone Methylation by NUE, a Novel Nuclear Effector of the Intracellular Pathogen Chlamydia trachomatis. PLOS Pathogens 2010, 6(7):e1000995.

18. Simon JM, Giresi PG, Davis IJ, Lieb JD: Using formaldehyde-assisted isolation of regulatory elements (FAIRE) to isolate active regulatory DNA. Nat Protoc 2012, 7(2):256–267.

19. Giresi PG, Kim J, McDaniell RM, Iyer VR, Lieb JD: FAIRE (Formaldehyde-Assisted Isolation of Regulatory Elements) isolates active regulatory elements from human chromatin. Genome Research 2007, 17(6):877–885.

20. Tan C, Hsia R-c, Shou H, Haggerty CL, Ness RB, Gaydos CA, Dean D, Scurlock AM, Wilson DP, Bavoil PM: Chlamydia trachomatis-infected patients display variable antibody profiles against the nine-member polymorphic membrane protein family. Infection and immunity 2009, 77(8):3218–3226.

21. Bolger AM, Lohse M, Usadel B: Trimmomatic: a flexible trimmer for Illumina sequence data. Bioinformatics (Oxford, England) 2014, 30(15):2114–2120.

22. FastQC: A Quality Control tool for High Throughput Sequence Data [http://www.bioinformatics.babraham.ac.uk/projects/fastqc/]

23. Langmead B, Salzberg SL: Fast gapped-read alignment with Bowtie 2. Nat Methods 2012, 9.

24. Wysoker A, Tibbetts K, Fennell T: Picard tools. http://picardsourceforgenet 2017.

25. Ramírez F, Dündar F, Diehl S, Grüning BA, Manke T: deepTools: a flexible platform for exploring deep-sequencing data. Nucleic Acids Research 2014, 42(Web Server issue):W187–W191.

26. Zhang Y, Liu T, Meyer CA, Eeckhoute J, Johnson DS, Bernstein BE, Nusbaum C, Myers RM, Brown M, Li W et al: Model-based Analysis of ChIP-Seq (MACS). Genome Biology 2008, 9(9):R137.

27. A comprehensive collection of signal artifact blacklist regions in the human genome. ENCODE. [hg19-blacklist-README.pdf] [http://mitra.stanford.edu/kundaje/akundaje/release/blacklists/hg38-human/]

28. DiffBind: Differential Binding Analysis of ChIP-Seq Peak Data. Bioconductor. [http://bioconductor.org/packages/release/bioc/html/DiffBind.html.]

29. Heinz S, Benner C, Spann N, Bertolino E, Lin YC, Laslo P, Cheng JX, Murre C, Singh H, Glass CK: Simple combinations of lineage-determining transcription factors prime cis-regulatory elements required for macrophage and B cell identities. Molecular cell 2010, 38(4):576–589.

30. Gao T, He B, Liu S, Zhu H, Tan K, Qian J: EnhancerAtlas: a resource for enhancer annotation and analysis in 105 human cell/tissue types. Bioinformatics (Oxford, England) 2016, 32(23):3543–3551.

31. Khan A, Zhang X: dbSUPER: a database of super-enhancers in mouse and human genome. Nucleic Acids Res 2016, 44(D1):D164–171.

32. Madani Tonekaboni SA, Mazrooei P, Kofia V, Haibe-Kains B, Lupien M: CREAM: Clustering of genomic REgions Analysis Method. bioRxiv 2018.

33. Khan A, Fornes O, Stigliani A, Gheorghe M, Castro-Mondragon JA, van der Lee R, Bessy A, Cheneby J, Kulkarni SR, Tan G et al: JASPAR 2018: update of the open-access database of transcription factor binding profiles and its web framework. Nucleic Acids Res 2018, 46(D1):D260–d266.

34. Gupta S, Stamatoyannopoulos JA, Bailey TL, Noble WS: Quantifying similarity between motifs. Genome Biology 2007, 8(2):R24–R24.

35. Gaulton KJ, Nammo T, Pasquali L, Simon JM, Giresi PG, Fogarty MP, Panhuis TM, Mieczkowski P, Secchi A, Bosco D et al: A map of open chromatin in human pancreatic islets. Nature genetics 2010, 42(3):255–259.

36. He Y, Carrillo JA, Luo J, Ding Y, Tian F, Davidson I, Song J: Genome-wide mapping of DNase I hypersensitive sites and association analysis with gene expression in MSB1 cells. Frontiers in genetics 2014, 5:308–308.

37. Gregory TR: Synergy between sequence and size in large-scale genomics. Nature reviews Genetics 2005, 6(9):699–708.

38. Ladomersky E, Khan A, Shanbhag V, Cavet JS, Chan J, Weisman GA, Petris MJ: Host and Pathogen Copper-Transporting P-Type ATPases Function Antagonistically during Salmonella Infection. Infection and immunity 2017, 85(9):e00351–00317.

39. Hodgkinson V, Petris MJ: Copper homeostasis at the host-pathogen interface. J Biol Chem 2012, 287(17):13549–13555.

40. Parnas O, Jovanovic M, Eisenhaure Thomas M, Herbst Rebecca H, Dixit A, Ye Chun J, Przybylski D, Platt Randall J, Tirosh I, Sanjana Neville E et al: A Genome-wide CRISPR Screen in Primary Immune Cells to Dissect Regulatory Networks. Cell 2015, 162(3):675–686.

41. Seaman MNJ: The retromer complex – endosomal protein recycling and beyond. Journal of cell science 2012, 125(20):4693–4702.

42. Elwell C, Engel J: Emerging Role of Retromer in Modulating Pathogen Growth. Trends in microbiology 2018, 26(9):769–780.

43. Paul B, Kim HS, Kerr MC, Huston WM, Teasdale RD, Collins BM: Structural basis for the hijacking of endosomal sorting nexin proteins by Chlamydia trachomatis. eLife 2017, 6.

44. Xu J, Zhang L, Ye Y, Shan Y, Wan C, Wang J, Pei D, Shu X, Liu J: SNX16 Regulates the Recycling of E-Cadherin through a Unique Mechanism of Coordinated Membrane and Cargo Binding. Structure (London, England : 1993) 2017, 25(8):1251–1263.e1255.

45. Schneider MR, Kolligs FT: E-cadherin’s role in development, tissue homeostasis and disease: Insights from mouse models: Tissue-specific inactivation of the adhesion protein E-cadherin in mice reveals its functions in health and disease. BioEssays : news and reviews in molecular, cellular and developmental biology 2015, 37(3):294–304.

46. Rajic J, Inic-Kanada A, Stein E, Dinic S, Schuerer N, Uskokovic A, Ghasemian E, Mihailovic M, Vidakovic M, Grdovic N et al: Chlamydia trachomatis Infection Is Associated with E-Cadherin Promoter Methylation, Downregulation of E-Cadherin Expression, and Increased Expression of Fibronectin and alpha-SMA-Implications for Epithelial-Mesenchymal Transition. Frontiers in cellular and infection microbiology 2017, 7:253.

47. Boehm M, Simson D, Escher U, Schmidt AM, Bereswill S, Tegtmeyer N, Backert S, Heimesaat MM: Function of Serine Protease HtrA in the Lifecycle of the Foodborne Pathogen Campylobacter jejuni. European journal of microbiology & immunology 2018, 8(3):70–77.

48. Backert S, Schmidt TP, Harrer A, Wessler S: Exploiting the Gastric Epithelial Barrier: Helicobacter pylori’s Attack on Tight and Adherens Junctions. Current topics in microbiology and immunology 2017, 400:195–226.

49. Wu X, Lei L, Gong S, Chen D, Flores R, Zhong G: The chlamydial periplasmic stress response serine protease cHtrA is secreted into host cell cytosol. BMC microbiology 2011, 11:87–87.

50. Gloeckl S, Ong VA, Patel P, Tyndall JDA, Timms P, Beagley KW, Allan JA, Armitage CW, Turnbull L, Whitchurch CB et al: Identification of a serine protease inhibitor which causes inclusion vacuole reduction and is lethal to Chlamydia trachomatis. Molecular microbiology 2013, 89(4):676–689.

51. Zhou Y, Zhu Y: Diversity of bacterial manipulation of the host ubiquitin pathways. Cellular microbiology 2015, 17(1):26–34.

52. Manzanillo PS, Ayres JS, Watson RO, Collins AC, Souza G, Rae CS, Schneider DS, Nakamura K, Shiloh MU, Cox JS: The ubiquitin ligase parkin mediates resistance to intracellular pathogens. Nature 2013, 501:512.

53. Haldar AK, Piro AS, Finethy R, Espenschied ST, Brown HE, Giebel AM, Frickel E-M, Nelson DE, Coers J: Chlamydia trachomatis Is Resistant to Inclusion Ubiquitination and Associated Host Defense in Gamma Interferon-Primed Human Epithelial Cells. mBio 2016, 7(6):e01417–01416.

54. Misaghi S, Balsara ZR, Catic A, Spooner E, Ploegh HL, Starnbach MN: Chlamydia trachomatis-derived deubiquitinating enzymes in mammalian cells during infection. Molecular microbiology 2006, 61(1):142–150.

55. Cocchiaro JL, Kumar Y, Fischer ER, Hackstadt T, Valdivia RH: Cytoplasmic lipid droplets are translocated into the lumen of the *Chlamydia trachomatis* parasitophorous vacuole. Proceedings of the National Academy of Sciences 2008, 105(27):9379–9384.

56. Guo B, Godzik A, Reed JC: Bcl-G, a novel pro-apoptotic member of the Bcl-2 family. J Biol Chem 2001, 276(4):2780–2785.

57. Baer M, Nilsen TW, Costigan C, Altman S: Structure and transcription of a human gene for H1 RNA, the RNA component of human RNase P. Nucleic Acids Res 1990, 18(1):97–103.

58. Egloff S, Studniarek C, Kiss T: 7SK small nuclear RNA, a multifunctional transcriptional regulatory RNA with gene-specific features. Transcription 2018, 9(2):95–101.

59. Ullu E, Weiner AM: Human genes and pseudogenes for the 7SL RNA component of signal recognition particle. The EMBO journal 1984, 3(13):3303–3310.

60. Hermanns P, Bertuch AA, Bertin TK, Dawson B, Schmitt ME, Shaw C, Zabel B, Lee B: Consequences of mutations in the non-coding RMRP RNA in cartilage-hair hypoplasia. Human molecular genetics 2005, 14(23):3723–3740.

61. Ho J, Moyes DL, Tavassoli M, Naglik JR: The Role of ErbB Receptors in Infection. Trends in microbiology 2017, 25(11):942–952.

62. Mölleken K, Becker E, Hegemann JH: The Chlamydia pneumoniae Invasin Protein Pmp21 Recruits the EGF Receptor for Host Cell Entry. PLOS Pathogens 2013, 9(4):e1003325.

63. Paes W, Dowle A, Coldwell J, Leech A, Ganderton T, Brzozowski A: The Chlamydia trachomatis PmpD adhesin forms higher order structures through disulphide-mediated covalent interactions. PLoS One 2018, 13(6):e0198662.

64. Patel AL, Chen X, Wood ST, Stuart ES, Arcaro KF, Molina DP, Petrovic S, Furdui CM, Tsang AW: Activation of epidermal growth factor receptor is required for Chlamydia trachomatis development. BMC microbiology 2014, 14:277.

65. Igietseme JU, Omosun Y, Nagy T, Stuchlik O, Reed MS, He Q, Partin J, Joseph K, Ellerson D, George Z et al: Molecular Pathogenesis of Chlamydia Disease Complications: Epithelial-Mesenchymal Transition and Fibrosis. Infect Immun 2018, 86(1).

66. van Ooij C, Kalman L, van I, Nishijima M, Hanada K, Mostov K, Engel JN: Host cell-derived sphingolipids are required for the intracellular growth of Chlamydia trachomatis. Cellular microbiology 2000, 2(6):627–637.

67. Elwell CA, Engel JN: Lipid acquisition by intracellular Chlamydiae. Cellular microbiology 2012, 14(7):1010–1018.

68. Miyazawa K, Miyazono K: Regulation of TGF-beta Family Signaling by Inhibitory Smads. Cold Spring Harbor perspectives in biology 2017, 9(3).

69. Takimoto T, Wakabayashi Y, Sekiya T, Inoue N, Morita R, Ichiyama K, Takahashi R, Asakawa M, Muto G, Mori T et al: Smad2 and Smad3 are redundantly essential for the TGF-beta-mediated regulation of regulatory T plasticity and Th1 development. Journal of immunology (Baltimore, Md : 1950) 2010, 185(2):842–855.

70. Attisano L, Tuen Lee-Hoeflich S: The Smads. Genome Biology 2001, 2(8):reviews3010.3011.

71. Hirokawa N, Noda Y, Tanaka Y, Niwa S: Kinesin superfamily motor proteins and intracellular transport. Nature Reviews Molecular Cell Biology 2009, 10:682.

72. Dumoux M, Menny A, Delacour D, Hayward RD: A Chlamydia effector recruits CEP170 to reprogram host microtubule organization. Journal of cell science 2015, 128(18):3420–3434.

73. Song L, Zhang Z, Grasfeder LL, Boyle AP, Giresi PG, Lee B-K, Sheffield NC, Gräf S, Huss M, Keefe D et al: Open chromatin defined by DNaseI and FAIRE identifies regulatory elements that shape cell-type identity. Genome research 2011, 21(10):1757–1767.

74. Lutter EI, Barger AC, Nair V, Hackstadt T: Chlamydia trachomatis inclusion membrane protein CT228 recruits elements of the myosin phosphatase pathway to regulate release mechanisms. Cell reports 2013, 3(6):1921–1931.

75. Grieshaber SS, Grieshaber NA, Hackstadt T: Chlamydia trachomatis uses host cell dynein to traffic to the microtubule-organizing center in a p50 dynamitin-independent process. Journal of cell science 2003, 116(Pt 18):3793–3802.

76. Lodish H BA, Zipursky SL, et al: Cilia and Flagella: Structure and Movement. In: Molecular Cell Biology. 4th edition edn: New York: W. H. Freeman; 2000.

77. Topham MK, Prescott SM: Mammalian diacylglycerol kinases, a family of lipid kinases with signaling functions. J Biol Chem 1999, 274(17):11447–11450.

78. Yao J, Cherian PT, Frank MW, Rock CO: Chlamydia trachomatis Relies on Autonomous Phospholipid Synthesis for Membrane Biogenesis. The Journal of biological chemistry 2015, 290(31):18874–18888.

79. Di Paolo Nelson C, Doronin K, Baldwin Lisa K, Papayannopoulou T, Shayakhmetov Dmitry M: The Transcription Factor IRF3 Triggers “Defensive Suicide” Necrosis in Response to Viral and Bacterial Pathogens. Cell Reports 2013, 3(6):1840–1846.

80. Gyorke CE, Nagarajan U: Interferon-Independent Protection by Interferon Regulatory Factor 3. The Journal of Immunology 2018, 200(1 Supplement):114.125–114.125.

81. Sixt BS, Bastidas RJ, Finethy R, Baxter RM, Carpenter VK, Kroemer G, Coers J, Valdivia RH: The Chlamydia trachomatis Inclusion Membrane Protein CpoS Counteracts STING-Mediated Cellular Surveillance and Suicide Programs. Cell Host Microbe 2017, 21(1):113–121.

82. Tan NY, Khachigian LM: Sp1 Phosphorylation and Its Regulation of Gene Transcription. Molecular and Cellular Biology 2009, 29(10):2483–2488.

83. Deniaud E, Baguet J, Chalard R, Blanquier B, Brinza L, Meunier J, Michallet M-C, Laugraud A, Ah-Soon C, Wierinckx A et al: Overexpression of Transcription Factor Sp1 Leads to Gene Expression Perturbations and Cell Cycle Inhibition. PLOS ONE 2009, 4(9):e7035.

84. Simmen RCM, Heard ME, Simmen AM, Montales MTM, Marji M, Scanlon S, Pabona JMP: The Krüppel-like factors in female reproductive system pathologies. Journal of molecular endocrinology 2015, 54(2):R89–R101.

85. Simmen RC, Heard ME, Simmen AM, Montales MT, Marji M, Scanlon S, Pabona JM: The Kruppel-like factors in female reproductive system pathologies. Journal of molecular endocrinology 2015, 54(2):R89–r101.

86. Dong JT, Chen C: Essential role of KLF5 transcription factor in cell proliferation and differentiation and its implications for human diseases. Cellular and molecular life sciences: CMLS 2009, 66(16):2691–2706.

87. Bieker JJ: Kruppel-like factors: three fingers in many pies. J Biol Chem 2001, 276(37):34355–34358.

88. Chiambaretta F, Nakamura H, De Graeve F, Sakai H, Marceau G, Maruyama Y, Rigal D, Dastugue B, Sugar J, Yue BY et al: Kruppel-like factor 6 (KLF6) affects the promoter activity of the alpha1-proteinase inhibitor gene. Investigative ophthalmology & visual science 2006, 47(2):582–590.

89. Swamynathan SK: Krüppel-like factors: three fingers in control. Human genomics 2010, 4(4):263–270.

90. Sun J, Wang B, Liu Y, Zhang L, Ma A, Yang Z, Ji Y, Liu Y: Transcription factor KLF9 suppresses the growth of hepatocellular carcinoma cells in vivo and positively regulates p53 expression. Cancer letters 2014, 355(1):25–33.

91. Subramaniam M, Hawse JR, Rajamannan NM, Ingle JN, Spelsberg TC: Functional role of KLF10 in multiple disease processes. BioFactors (Oxford, England) 2010, 36(1):8–18.

92. Truty MJ, Lomberk G, Fernandez-Zapico ME, Urrutia R: Silencing of the transforming growth factor-beta (TGFbeta) receptor II by Kruppel-like factor 14 underscores the importance of a negative feedback mechanism in TGFbeta signaling. J Biol Chem 2009, 284(10):6291–6300.

93. Chasman DI, Paré G, Mora S, Hopewell JC, Peloso G, Clarke R, Cupples LA, Hamsten A, Kathiresan S, Mälarstig A et al: Forty-three loci associated with plasma lipoprotein size, concentration, and cholesterol content in genome-wide analysis. PLoS genetics 2009, 5(11):e1000730–e1000730.

94. Sarmento OF, Svingen PA, Xiong Y, Xavier RJ, McGovern D, Smyrk TC, Papadakis KA, Urrutia RA, Faubion WA: A Novel Role for Kruppel-like Factor 14 (KLF14) in T-Regulatory Cell Differentiation. Cellular and Molecular Gastroenterology and Hepatology 2015, 1(2):188–202.e184.

95. de Ruijter AJ, van Gennip AH, Caron HN, Kemp S, van Kuilenburg AB : Histone deacetylases (HDACs): characterization of the classical HDAC family. The Biochemical journal 2003, 370(Pt 3):737–749.

