## Supplementary figures and images for "Chromatin accessibility dynamics of *Chlamydia*-infected epithelial cells"

### Additional File 1.docx

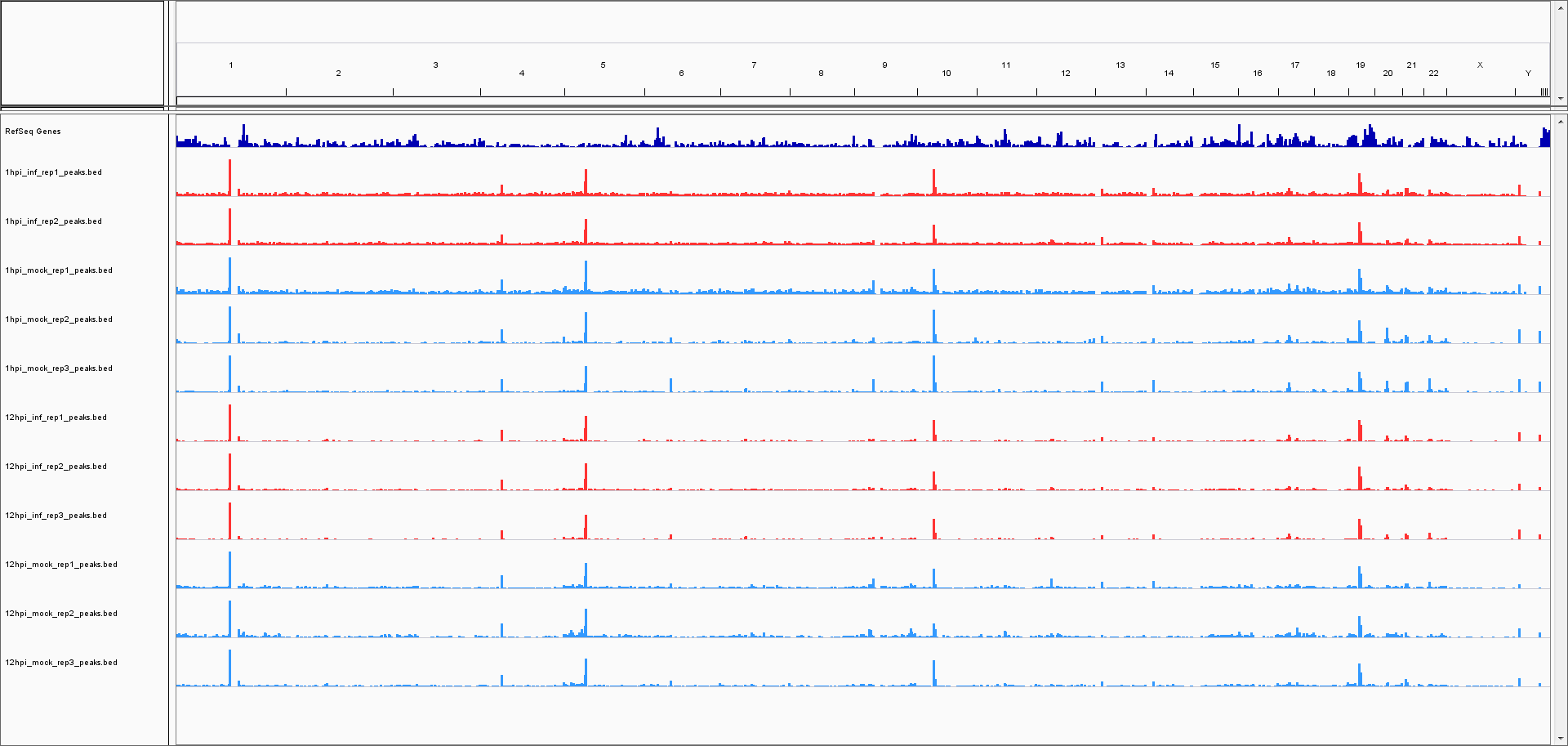

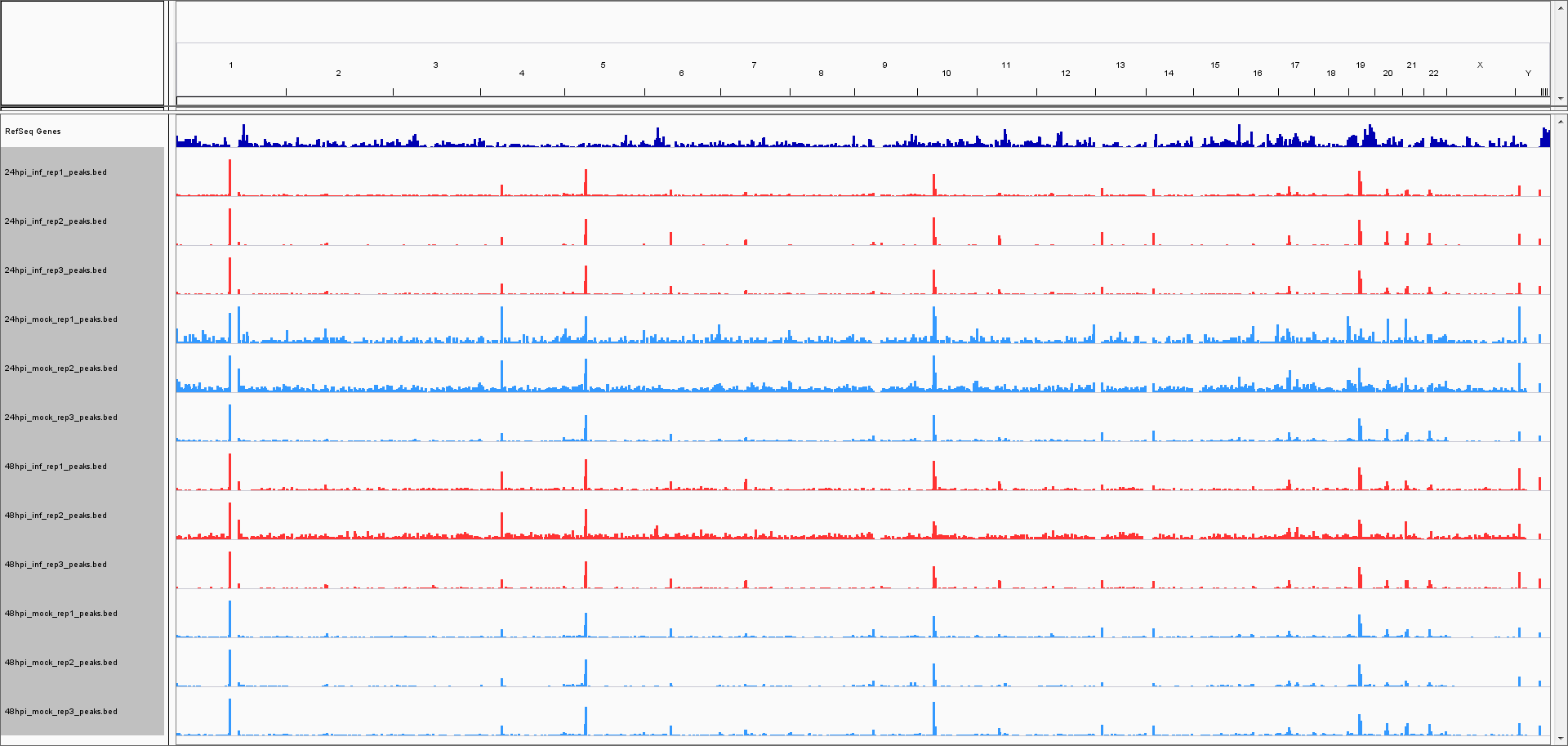
