## Additional File 3.docx for "Chromatin accessibility dynamics of *Chlamydia*-infected epithelial cells"

**Enrichment of Super-enhancers**

| **Time** | **Category** | **Gene** | **Full name** | **Description** |
| --- | --- | --- | --- | --- |
| 24 | Signalling | MSN | Moesin (Membrane-Organizing Extension Spike Protein) | Belongs to the ERM family of proteins (ezrin/radixin/moesin) acting as signalling molecules inducing cytoskeletal organisation and remodelling^1^. |
| 48 | Cell structure and cell Signalling | FLNB | Filamin B | Helps build the cytoskeleton through a network of protein filaments, giving structure to cells. Filamin B binds to actin which supports a range of functions including cell signalling directing the cytoskeletal changes during growth and development^2^ |
| 48 | Signalling | PTP4A2 | Protein Tyrosine Phosphatase Type IVA, Member 2 | PTP’s are cell signalling molecules. Involved with plasmic and endosomal membranes^3^. Also shown to stimulate progression during mitosis from G1 to S phase^4^ |
| 48 | Cell growth | KLF5 | Kruppel Like Factor 5 | Zinc finger (transcription factor) that is a transcriptional activator, binding to promoters of target genes^5^.  Linked to cell growth and depending on tissue and conditions can stimulate or supress growth^6^. |
| 48 | Innate immune response | IER3 | Immediate Early Response 3 | Induced from a wide arrange of stimuli including growth factors, cytokines, viral infection and other stressors^7^. Main function is involved in protection of the cell. |
| 48 | DNA-based binding | RREB1 | Ras Responsive Element Binding Protein 1 | Zinc finger (TF) that increases transcription and binds to promoter regions of target genes (RAS-responsive elements - RRE)^8^ |
| 48 | Protein-binding | ATXN10 | Ataxin 10 | May be involved in intracellular glycosylation homeostasis^9^. |

**References**

1	Abiatari, I. *et al.* Moesin-dependent cytoskeleton remodelling is associated with an anaplastic phenotype of pancreatic cancer. *J Cell Mol Med* **14**, 1166-1179, doi:10.1111/j.1582-4934.2009.00772.x (2010).

2	NIH. *FLNB gene*, <<https://ghr.nlm.nih.gov/gene/FLNB>> (2019).

3	NCBI. *PTP4A2 protein tyrosine phosphatase 4A2 - Homo Sapiens*, <<https://www.ncbi.nlm.nih.gov/gene/8073#general-gene-info>> (2019).

4	Uniprot. *UniProtKB - Q12974 (TP4A2_HUMAN)*, <<https://www.uniprot.org/uniprot/Q12974#function>> (2019).

5	NCBI. *KLF5 Kruppel like factor 5 - Homo sapiens (human)*, <<https://www.ncbi.nlm.nih.gov/gene/688>> (2019).

6	Diakiw, S. M., D'Andrea, R. J. & Brown, A. L. The double life of KLF5: Opposing roles in regulation of gene-expression, cellular function, and transformation. *IUBMB Life* **65**, 999-1011, doi:10.1002/iub.1233 (2013).

7	Arlt, A. & Schafer, H. Role of the immediate early response 3 (IER3) gene in cellular stress response, inflammation and tumorigenesis. *European journal of cell biology* **90**, 545-552, doi:10.1016/j.ejcb.2010.10.002 (2011).

8	NCBI. *RREB1 ras responsive element binding protein 1 - Homo sapiens (human)*, <<https://www.ncbi.nlm.nih.gov/gene/6239>> (2019).

9	Uniprot. *UniProtKB - Q9UBB4 (ATX10_HUMAN)*, <<https://www.uniprot.org/uniprot/Q9UBB4#function>> (2019).
